# Ca^2+^-activated sphingomyelin scrambling and turnover mediate ESCRT-independent lysosomal repair

**DOI:** 10.1101/2021.03.12.435146

**Authors:** Patrick Niekamp, Tolulope Sokoya, Laura Vittadello, Yongqiang Deng, Yeongho Kim, Angelika Hilderink, Mirco Imlau, Christopher J. Clarke, Christopher G. Burd, Joost C. M. Holthuis

## Abstract

Lysosomes are vital organelles vulnerable to injuries from diverse materials. Failure to repair or sequester damaged lysosomes poses a threat to cell viability. Here we report that cells exploit a sphingomyelin-based lysosomal repair pathway that operates independently of ESCRT to reverse potentially lethal membrane damage. Various conditions perturbing organelle integrity trigger a rapid calcium-activated scrambling and cytosolic exposure of sphingomyelin. Subsequent metabolic conversion of sphingomyelin by neutral sphingomyelinases on the cytosolic surface of injured lysosomes promotes their repair, also when ESCRT function is compromised. Conversely, blocking turnover of cytosolic sphingomyelin renders cells more sensitive to lysosome-damaging drugs. Our data indicate that calcium-activated scramblases, sphingomyelin, and neutral sphingomyelinases are core components of a previously unrecognized membrane restoration pathway by which cells preserve the functional integrity of lysosomes.

---

Lysosomes are essential cellular organelles involved in the degradation of macromolecules, pathogen killing and metabolic signaling. To perform these vital tasks, lysosomes contain high concentrations of acid hydrolases, protons and calcium. Conversely, lysosomal damage caused by incoming pathogens, amphiphilic drugs or sharp crystals can have deleterious consequences, including cell death^1^. To avoid spilling of harmful lysosomal contents into the cytosol, injured lysosomes are marked for degradation by a specialized form of autophagy, known as lysophagy. This process is initiated by recruitment of cytosolic galectins and glycoprotein-specific ubiquitin ligases to abnormally exposed luminal glycans at the lesion site, resulting in engulfment of the damaged lysosome by autophagic membranes^2,3^. While lysophagy is a slow process in which the disrupted organelle is ultimately sacrificed, recent work revealed an important role of the Endosomal Sorting Complex Required for Transport (ESCRT) machinery in repairing small perforations in lysosomes to allow their escape from autophagic degradation^4,5^. ESCRT proteins are organized in functionally distinct complexes that drive an inverse membrane remodeling during various cellular processes, including cytokinetic abscission, vesicle biogenesis inside multivesicular endosomes, and viral budding, in addition to membrane repair^6–8^. All these processes require ESCRT-III proteins that form filaments within membrane invaginations and cooperate with the ATPase VSP4 to catalyze membrane constriction and fission away from the cytosol^9^.

Activation of ESCRT enables cells to prevent potentially lethal consequences of minor perturbations in lysosomal integrity via a mechanism that seems to sense more subtle membrane injuries than galectins. However, the recruitment signal that triggers ESCRT-III assembly at sites of lysosomal damage has not been established with certainty. While detection of Ca^2+^ leakage out of the injured lysosome by ALIX and its Ca^2+^-binding partner ALG2 has been proposed as a mechanism^4^, other studies were unable to confirm a requirement for Ca^2+^ and found that the ESCRT-I subunit TSG101 is more important than ALIX for mediating ESCRT-III recruitment in this context^5,10^. Thus, ESCRT assembly on damaged organelles may involve hitherto uncharacterized cues^6^.

While glycans reside exclusively on the non-cytosolic surface of lysosomes and the plasma membrane, also certain lipids display strict asymmetric distributions across organellar bilayers. For instance, sphingomyelin (SM) is highly enriched in the exoplasmic leaflet of the plasma membrane whereas phosphatidylserine (PS) is primarily located in the cytosolic leaflet^11–13^. Translocation of PS to the outer leaflet during apoptosis marks dying cells and leads to their timely removal^14^. Application of a GFP-tagged version of lysenin, a SM-specific toxin from the earthworm *Eisenia fetida*^15^, revealed that SM becomes exposed to the cytosol upon endomembrane damage caused by Gram-negative pathogens like *Shigella flexneri* or *Salmonella typhimurium*^16^. Breakout of these pathogens from the host vacuole into the cytosol follows a multi-step process, in which cytosolic SM exposure detected by the lysenin-based reporter invariably preceded glycan exposure, catastrophic membrane damage, and cytosolic entry of the bacteria. This raised the idea that the arrival of SM on the cytosolic surface of bacteria-containing vacuoles provides an early warning signal to alert cells of an imminent break down of organellar integrity^16^. How SM transfer across the bilayer of injured organelles is initiated and whether this process is part of a mechanism that helps preserve the integrity of cellular organelles remain to be explored.

## RESULTS

### SM is readily exposed on the cytosolic surface of damaged organelles

To study how membrane damage triggers transbilayer movement of SM, we used an engineered version of the SM-binding pore-forming toxin, equinatoxin II (Eqt). Expression of EqtSM carrying a *N*-terminal signal sequence and *C*-terminal GFP tag previously enabled us to demonstrate sorting of native SM at the *trans*-Golgi network into a distinct class of secretory vesicles^17^. When expressed without signal sequence, EqtSM displayed a diffuse distribution throughout the cytosol and nucleus of HeLa cells (Fig. 1A, 0 min time point). Occasionally, the cytosolic reporter gave rise to a few small intracellular puncta, which may originate from a minor tendency of Eqt to aggregate upon overexpression or SM exposure following spontaneous membrane damage. However, in cells treated with L-leucyl-L-leucine O-methyl ester (LLOMe), a lysosomotropic compound commonly used to disrupt lysosome integrity^18^, EqtSM underwent a massive accumulation in numerous puncta distributed throughout the cell within minutes after drug addition (Fig. 1A, B). In contrast, LLOMe failed to trigger recruitment of a SM binding-deficient Eqt mutant, EqtSol. LLOMe-induced mobilization of EqtSM was virtually abolished upon genetic ablation of the SM synthases SMS1 and SMS2 (SMS-KO; Fig. 1A, B; Fig. S1A). While SMS removal essentially abolished the cellular SM pool, it did not impair the ability of LLOMe to disrupt lysosomal integrity (Fig. S2). Collectively, these data indicate that EqtSM faithfully reports cytosolic exposure of SM by LLOMe-damaged lysosomes.

**Fig. 1.**
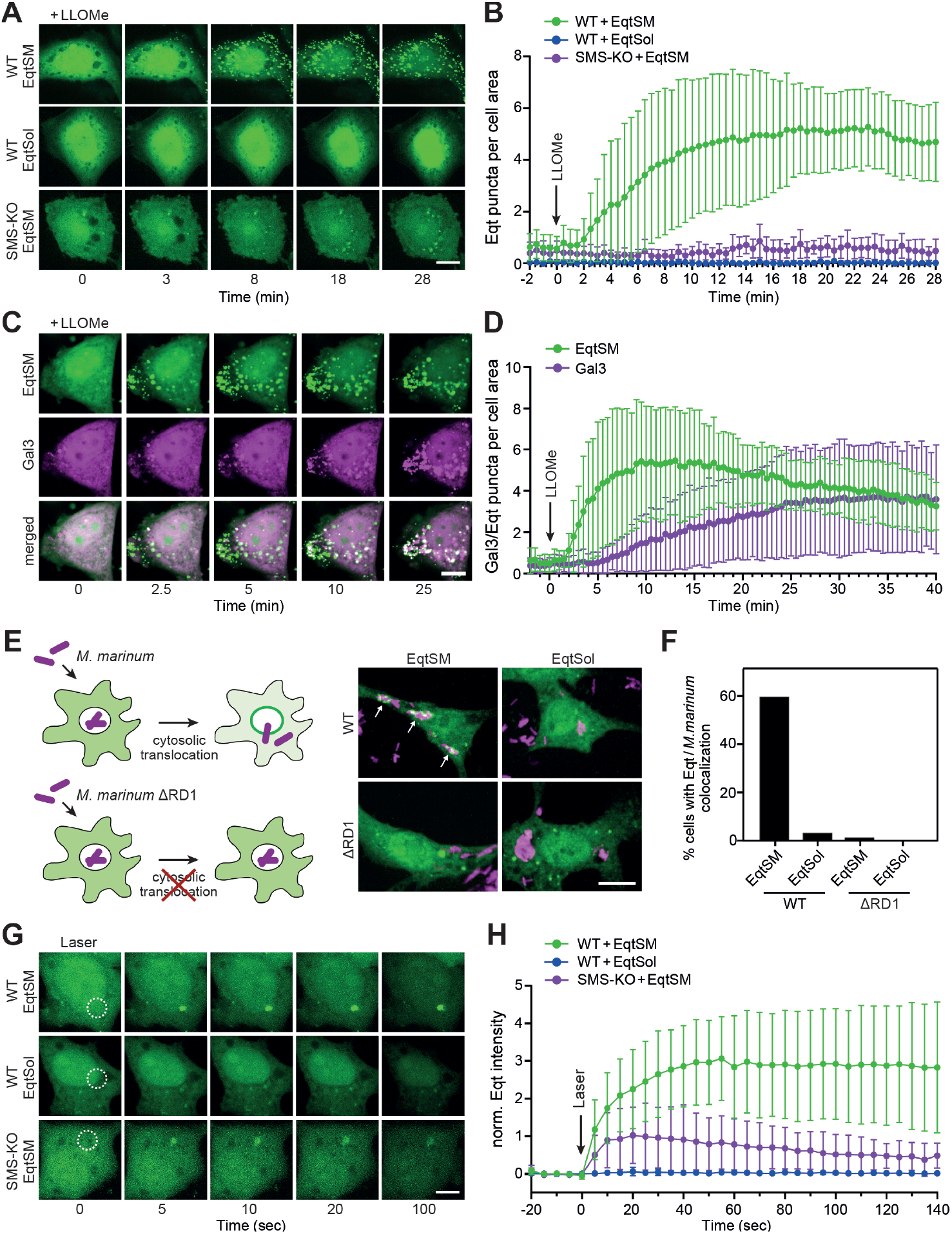
Cytosolic EqtSM readily binds organelles injured by chemicals, pathogens or light. (**A**) Time-lapse fluorescence images of wildtype (WT) or SMS-KO HeLa cells expressing GFP-tagged EqtSM or EqtSol and treated with 1 mM LLOMe for the indicated time. (**B**) Time-course plotting Eqt-positive puncta per 100 μm^2^ cell area in cells treated as in (A). Data are means ± SD from ≥ 8 cells per condition. (**C**) Time-lapse fluorescence images of wildtype HeLa cells co-expressing GFP-tagged EqtSM (*green*) and mCherry-tagged Gal3 (*magenta*) treated with 1 mM LLOMe for the indicated time. (**D**) Time-course plotting Eqt- and Gal3-positive puncta per 100 μm^2^ cell area in cells treated as in (C). Data are means ± SD from ≥ 17 cells per condition. (**E**) RAW264.7 cells expressing GFP-tagged EqtSM or EqtSol (*green*) were infected with mCherry-expressing wildtype (WT) or translocation-defective (ΔRD1) mutant strains of *Mycobacteria marinum* (*magenta*). Live-cell fluorescence micrographs were captured 2 h post-infection. (**F**) Percentage of cells treated as in (E) showing co-localization of mCherry-expressing *M. marinum* and EqtSM-positive puncta. Data are means from ≥ 30 cells per condition. (**G**) Time-lapse fluorescence images of wildtype (WT) or SMS-KO HeLa cells expressing GFP-tagged EqtSM or EqtSol and locally wounded by a brief pulse from a 2-photon laser. (**H**) Time-course plotting Eqt-associated fluorescence at the laser-induced wound site in cells treated as in (G). Data are means ± SD from ≥ 5 cells per condition. Scale bar, 10 μm.

To analyze the kinetics of EqtSM mobilization in relation to the degree of lysosomal damage, we next used cells co-expressing the SM reporter with mCherry-tagged Galectin-3 (Gal3), a cytosolic lectin with affinity for the complex glycans that reside on the non-cytosolic surface of lysosomes^19^. Upon LLOMe treatment, EqtSM readily accumulated in numerous puncta that also became gradually positive for Gal3 (Fig. 1C; Movie S1). Importantly, EqtSM was recruited to LLOMe-damaged lysosomes prior to Gal3, with a time difference of approximately 5 min (Fig. 1D). These findings are consistent with those reported by Ellison *et al.*^16^ and indicate that a break in SM asymmetry is an early marker of lysosomal damage that precedes a catastrophic breakdown of membrane integrity, when galectins gain access to the luminal glycans. Live-cell imaging of RAW246.7 macrophages invaded by the bacterial pathogen *Mycobacterium marinum* revealed strong recruitment of EqtSM to a subset of bacteria at 1-2 h post-infection (Fig. 1E, F; Movie S2). No EqtSM recruitment was observed in macrophages invaded by the non-pathogenic *M. marinum* strain ΔRD1, which fails to translocate to the cytosol and remains confined to the bacteria-containing phagosome due to a non-functional ESX-1 secretion system required for niche-breakage and pore-forming activity^20,21^.

To further challenge a fundamental link between transbilayer SM movement and membrane damage, we next analysed the EqtSM distribution in cells exposed to distinct modes of plasma membrane-wounding. To this end, we first incubated cells expressing EqtSM with the bilayer-destabilizing compound digitonin. We observed a redistribution of the cytosolic SM reporter to discrete puncta at the plasma membrane within minutes after transient exposure to the compound (Fig. S3A). Similar results were obtained when cells were incubated with the pore-forming bacterial toxin streptolysin O (SLO; Fig. S3B). To couple localized plasma membrane injuries to a fast imaging of downstream events, we next conducted laser-based plasma membrane wounding using a confocal scanning microscope equipped with a pulsed-laser. This revealed an ultra-fast (within 5 sec) and localized mobilization of EqtSM, but not EqtSol, to the laser-induced wound site (Fig. 1G, H; Movie S3). While EqtSM is readily recruited to the damaged membrane area in both wildtype and SMS-KO cells, the signal in the latter readily faded. This is in line with our finding that SMS-KO cells contain only residual amounts of SM (Fig. S2A). Thus, a breach in membrane integrity caused by pore-forming chemicals, bacterial toxins or a laser appears to be tightly linked to a rapid transbilayer movement of SM.

### Damage triggers SM translocation through calcium-activated scramblases

An influx of calcium from the extracellular environment or calcium-storing organelles has been recognized as a central trigger in the detection and repair of membrane injuries^22,23^. To address whether calcium plays a role in the damage-induced translocation of SM, we next analysed the distribution of cytosolic EqtSM in HeLa cells exposed to SLO in calcium-free medium supplemented with the calcium chelator EGTA. Removal of extracellular calcium greatly impaired SLO-induced formation of EqtSM-positive puncta (Fig. 2A, B). Moreover, treatment of cells with the calcium ionophore ionomycin triggered an accumulation of EqtSM in numerous puncta, but only when calcium was present in the medium (Fig. 2C, D). The plasma membrane of mammalian cells harbors a calcium-activated phospholipid scramblase, TMEM16F, which catalyzes phosphatidylserine (PS) externalization in response to elevated intracellular calcium^24^. We therefore wondered whether TMEM16F plays a role in SM scrambling at sites of plasma membrane damage. Strikingly, genetic ablation of TMEM16F abolished the calcium-dependent formation of EqtSM-positive puncta in both SLO- and ionomycin-treated cells (Fig. 2A-D, Fig. S4). This indicates that TMEM16F is responsible for damage-induced SM movement across the plasma membrane.

**Fig. 2.**
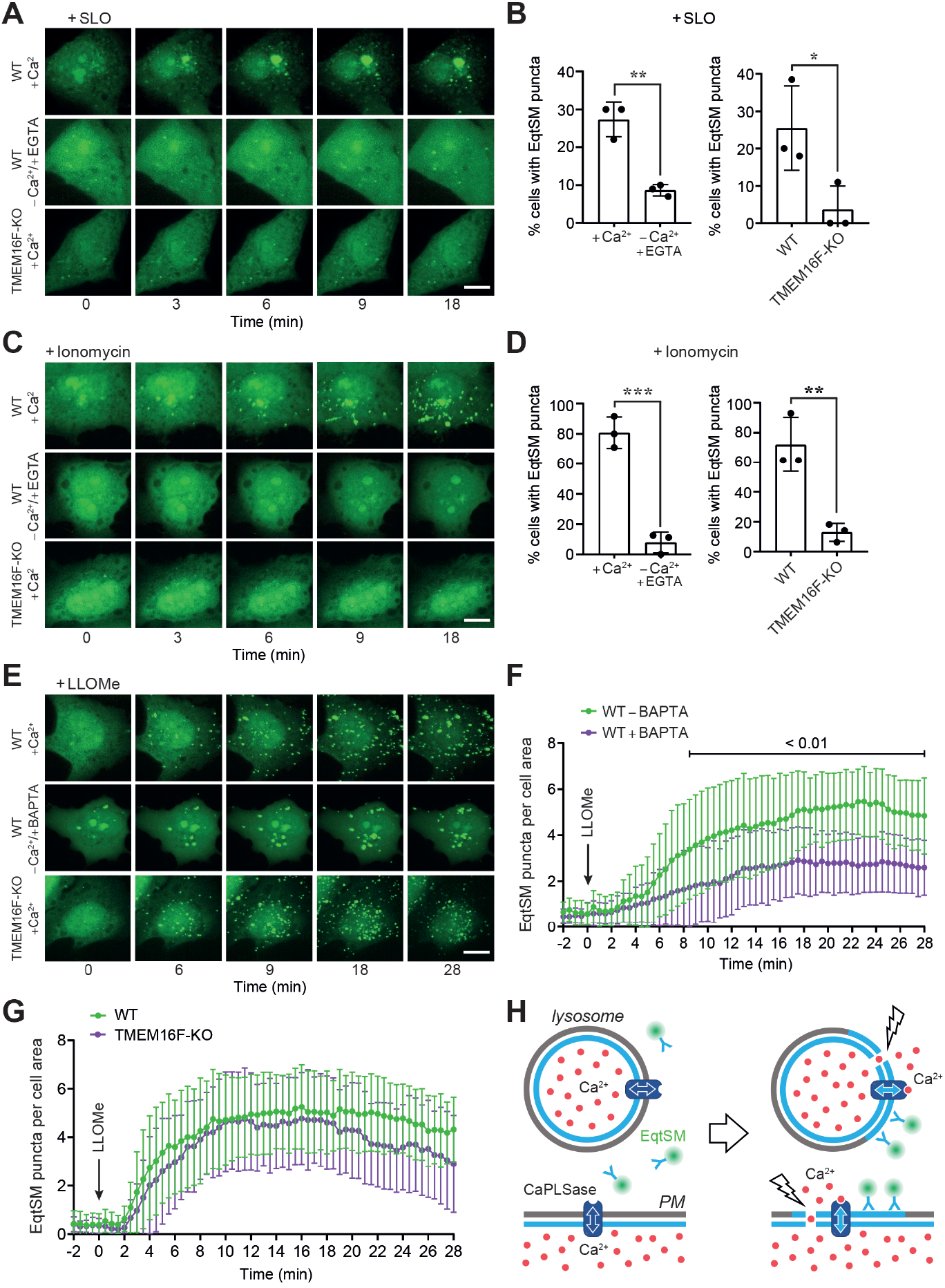
Damage-induced SM translocation is mediated by a calcium-activated lipid scramblase. (**A**) Time-lapse fluorescence images of wildtype (WT) or TMEM16F-KO HeLa cells expressing GFP-tagged EqtSM and treated with 1500 U/ml SLO for the indicated time in medium containing (+Ca^2+^) or lacking Ca^2+^ (-Ca^2+^/+EGTA). (**B**) Percentage of cells displaying EqtSM-positive puncta after 30 min of SLO treatment as in (A). Data are means ± SD from ≥ 24 cells per condition, *n* = 3. \**p* ≤ 0.05 and \*\**p* ≤ 0.01 by unpaired two-tailed t-test. (**C**) Time-lapse fluorescence images of wildtype (WT) or TMEM16F-KO HeLa cells expressing GFP-tagged EqtSM and treated with 5 μM ionomycin for the indicated time in medium containing (+Ca^2+^) or lacking Ca^2+^ (-Ca^2+^/ +EGTA). (**D**) Percentage of cells displaying EqtSM-positive puncta after 30 min of ionomycin treatment as in (A). Data are means ± SD from ≥ 24 cells per condition, *n* = 3. \*\**p* ≤ 0.01 and \*\*\**p* ≤ 0.001 by unpaired two-tailed t-test. (**E**) Time-lapse fluorescence images of wildtype (WT) or TMEM16F-KO HeLa cells expressing GFP-tagged EqtSM treated with 1 mM LLOMe after pre-incubation with or without BAPTA-AM (100 μM, 45 min). Scale bar, 10 μm. (**F**) Time-course plotting EqtSM-positive puncta per 100 μm^2^ cell area in wildtype (WT) cells treated as in (E). Data are means ± SD from ≥ 21 cells per condition, *n* = 3. Statistical significance was determined by unpaired two-tailed t-test. (**G**) Time-course plotting EqtSM-positive puncta per 100 μm^2^ cell area in wildtype or TMEM16F-KO HeLa cells treated with LLOMe in the absence of BAPTA-AM as in (F). Data are means ± SD from ≥ 21 cells per condition, *n* = 3. (**H**) Schematic illustration of how membrane damage triggers SM scrambling. PM, plasma membrane; CaPLSase, calcium-activated phospholipid scramblase.

Because lysosomes store calcium and rupturing them increases intracellular calcium, we next asked whether EqtSM recruitment to damaged lysosomes is similarly controlled by calcium-activated scramblases. Preloading cells with the lysosomal calcium chelator BAPTA-AM significantly impaired recruitment of the cytosolic SM reporter to LLOMe-injured lysosomes (Fig. 2E, F). In contrast, removal of TMEM16F did not affect the drug-induced lysosomal mobilization of the reporter (Fig. 2E, G). Based on these results, we conclude that the transbilayer movement of SM in response to membrane injuries is a calcium-dependent process, involving TMEM16F at the plasma membrane and presumably a related scramblase in lysosomes (Fig. 2H). The identity of the lysosomal scramblase remains to be established.

### SM-deficient cells are defective in lysosomal repair

We next investigated whether SM actively participates in the mechanism by which cells detect and repair damaged lysosomes. To this end, we took advantage of the ability of lysosomes to retain LysoTracker, a weak base that accumulates in the organelle’s acidic lumen and is fluorescent at low pH^25^. LysoTracker fluorescence is lost upon brief exposure of cells to the lysosomotropic compound glycyl-L-phenylalanine 2-naphtylamide (GPN) but returns after GPN is washed away (Fig. 3A, B). Like LLOMe, GPN induces a transient disruption of lysosomal integrity that is accompanied by an accumulation of EqtSM in numerous puncta (Fig. 3B, C; Movie S4). However, GPN is processed into metabolites thought to promote osmotic rupture^26^. As lysosomal pH critically relies on the structural integrity of the membrane, the loss and recovery of LysoTracker fluorescence in cells transiently exposed to GPN is consistent with lysosomal membrane disruption and restoration. Wildtype and SMS-KO cells acquired LysoTracker to a similar extent (Fig. S5), and also lost LysoTracker fluorescence with similar kinetics after addition of GPN (Fig. 3D, E), indicating comparable lysosome function. However, recovery of LysoTracker fluorescence after GPN removal was significantly delayed in the SMS-KO cells. Moreover, less LysoTracker-positive structures recovered in SMS-KO cells than in wildtype. In SMS-KO cells transduced with SMS1 under control of a doxycycline-inducible promotor, addition of doxycycline restored the capacity to produce SM (Fig. S1) and regain LysoTracker fluorescence to the same level as in wildtype cells after GPN is washed away (Fig. 3F, G). SMS removal also caused a significant rise in cell death in the presence of LLOMe (Fig. 3H). Collectively, this indicates that SM plays a critical role in the recovery of lysosomes from acute, potentially lethal damage.

**Fig. 3.**
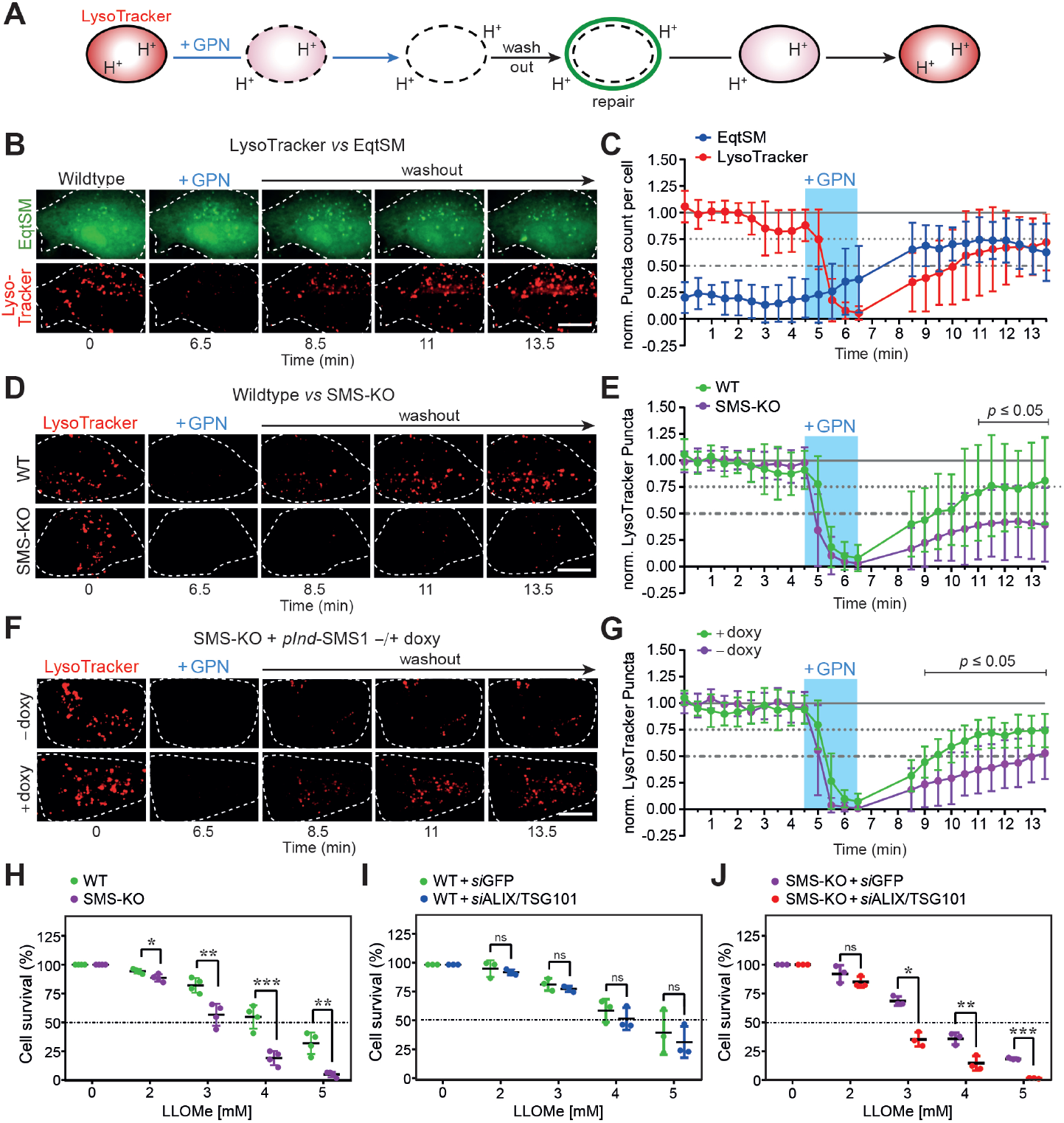
SM is critical for recovery of lysosomes from acute damage. (**A**) Schematic outline of lysosomal repair assay. Cells are incubated with LysoTracker to label functional lysosomes. Externally added GPN disrupts the lysosomal membrane, causing a proton efflux and loss of LysoTracker fluorescence. Upon GPN washout, LysoTracker fluorescence gradually recovers, providing a measure for lysosomal repair. (**B**) Time-lapse fluorescence images of LysoTracker-labeled and EqtSM-expressing HeLa cells during and after a 2 min-pulse of GPN (200 μM). Scale bar, 10 μm. (**C**) Time-course plotting LysoTracker- and EqtSM-positive puncta in cells treated as in (B), normalized to the initial number of puncta. Data are means ± SD from 18 cells, *n* = 4. (**D**) Time-lapse fluorescence images of LysoTracker-labeled wildtype (WT) and SMS-KO HeLa cells during and after a 2 min-pulse of GPN. Scale bar, 10 μm. (**E**) Time-course plotting LysoTracker-positive puncta in cells treated as in (B), normalized to the initial number of puncta. Data are means ± SD from ≥ 24 cells per condition, *n* = 4. Statistical significance was determined by unpaired two-tailed t-test. (**F**) Time-lapse fluorescence images of LysoTracker-labeled SMS-KO HeLa cells transduced with doxycycline-inducible SMS1 (*pInd*-SMS1) during and after a 2 min-pulse of GPN following 48 h pre-incubation in the absence (-doxy) or presence of 1 μg/ml doxycycline (+doxy). Scale bar, 10 μm. (**G**) Time-course plotting LysoTracker-positive puncta in cells treated as in (D), normalized to the initial number of puncta. Data are means ± SD from ≥ 17 cells per condition, *n* = 3. Statistical significance was determined by unpaired two-tailed t-test. (**H**) Survival rate of wildtype (WT) and SMS-KO HeLa cells after 5 h exposure to LLOMe at the indicated concentration. Data are means ± SD, *n* = 4. (**I**) Survival rate of wildtype (WT) HeLa cells pre-treated with siRNAs targeting GFP or ALIX and TSG101 for 72 h and then exposed for 5 h to LLOMe at the indicated concentration. Data are means ± SD, *n* = 3. (**J**) Survival rate of SMS-KO HeLa cells pre-treated with siRNAs targeting GFP or ALIX and TSG101 for 72 h and then exposed for 5 h to LLOMe at the indicated concentration. Data are means ± SD, *n* = 3. Statistical significance was determined by paired two-tailed t-test. *p*-values are presented where significant. \**p* ≤ 0.05, \*\**p* ≤ 0.01, \*\*\**p* ≤ 0.001.

### SM is dispensable for ESCRT recruitment to damaged lysosomes

Previous work revealed that the ESCRT machinery is readily recruited to damaged lysosomes and participates in their repair^4,5^. As ESCRT recruitment is at least partially dependent on a rise in intracellular calcium^4^, we wondered whether cytosolic SM exposure may be part of the mechanism by which ESCRT is mobilized to injured lysosomes. To address this idea, we first monitored the subcellular distribution of both EqtSM and CHMP4B, an ESCRT-III component necessary for all known mammalian ESCRT functions^6^, in LLOMe-treated cells. This revealed that mobilization of EqtSM to LLOMe-damaged lysosomes precedes CHMP4B recruitment with a time gap of about 60 seconds (Fig. 4A, B; Movie S5). We then asked whether SM is required for mobilizing this central ESCRT-III component. However, the rate and efficiency by which CHMP4B accumulated on LLOMe-damaged lysosomes in SMS-KO cells were indistinguishable from those in wildtype cells (Fig. 4C, D). On the other hand, SMS-KO cells displayed a prolonged CHMP4B retention on damaged lysosomes, a finding consistent with a critical role of SM in lysosomal repair. Moreover, siRNA-mediated depletion of ALIX and TSG101, two proteins essential for ESCRT-III recruitment to damaged lysosomes ^5^, further reduced the viability of LLOMe-treated SMS-KO cells while having only a minor impact on the survival of LLOMe-treated controls (Fig. 3I, J; Fig. S6). This indicates that cells are equipped with a SM-dependent lysosomal repair pathway that operates in parallel with the ESCRT pathway.

**Fig. 4.**
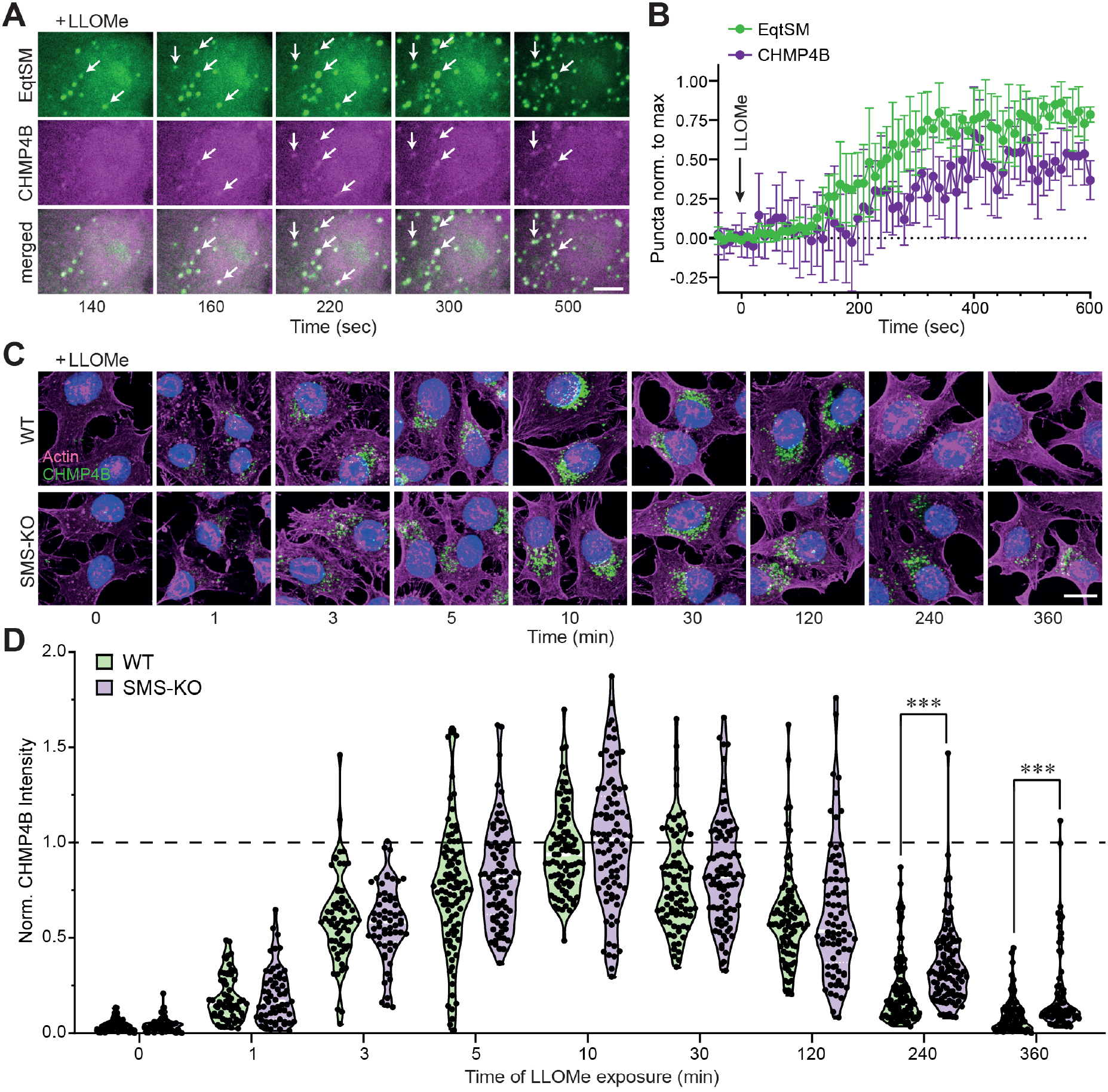
SM is dispensable for ESCRT recruitment to damaged lysosomes. (**A**) Time-lapse fluorescence images of wildtype HeLa cells co-expressing mKate-tagged EqtSM (*green*) and eGFP-tagged CHMP4B (*magenta*) and treated with 1 mM LLOMe for the indicated time. White arrows mark EqtSM-positive puncta that gradually accumulate CHMP4B. (**B**) Time-course plotting EqtSM- and CHMP4B-positive puncta normalized to maximum in cells treated as in (A). Data are means ± SD from 5 cells. (**C**) Fluorescence images of wildtype (WT) and SMS-KO HeLa cells treated with 1 mM LLOMe for the indicated time, fixed, and then stained with DAPI (*blue*) and antibodies against CHMP4B (*green*) and actin (*magenta*). (**D**) Time-course plotting normalized CHMP4B intensity in cells treated as in (C). More than 30 cells per condition were analyzed, *n* = 2. Statistical significance was determined by unpaired two-tailed t-test. *p*-values are presented where significant. \*\*\**p* ≤ 0.001.

### Hydrolysis of cytosolic SM enhances lysosomal repair in ESCRT-compromised cells

To investigate whether SM exposure on the cytosolic surface of damaged lysosomes is critical for their repair, we generated a construct in which a SMase from *Bacillus cereus* was fused to the cytosolic tail of the lysosome-associated membrane protein LAMP1 (Fig. 5A). The LAMP1-bSMase fusion protein co-localized with LAMP1-positive lysosomes and catalyzed the metabolic conversion of fluorescent NBD-SM into NBD-ceramide (Fig. S7). LAMP1 fused to a catalytically inactive bSMase mutant (D322A/H323A) served as control. Live cell imaging revealed that expression of active LAMP1-bSMase, but not its enzyme-dead counterpart, efficiently suppressed mobilization of cytosolic EqtSM to LLOMe-damaged lysosomes (Fig. 5B, C; Movies S6 and S7), indicating that the active fusion protein catalyzes a rapid and efficient hydrolysis of SM on the cytosolic surface of damaged lysosomes. To address whether metabolic turnover of cytosolic SM reduces or enhances the repair of damaged lysosomes, we next analyzed the impact of active or enzyme-dead LAMP1-bSMase on the ability of cells to regain LysoTracker fluorescence after transient exposure to GPN. We found that expression of active LAMP1-bSMase enhanced rather than diminished the recovery of LysoTracker fluorescence in GPN-treated cells (Fig. 5D). This effect became even more prominent when these experiments were carried out on cells treated with ALIX and TSG101-targeting siRNAs (Fig. 5E). Thus, a calcium-induced exposure and subsequent metabolic turnover of SM on the cytosolic surface of damaged lysosomes promotes their repair, even when ESCRT recruitment is blocked.

**Fig. 5.**
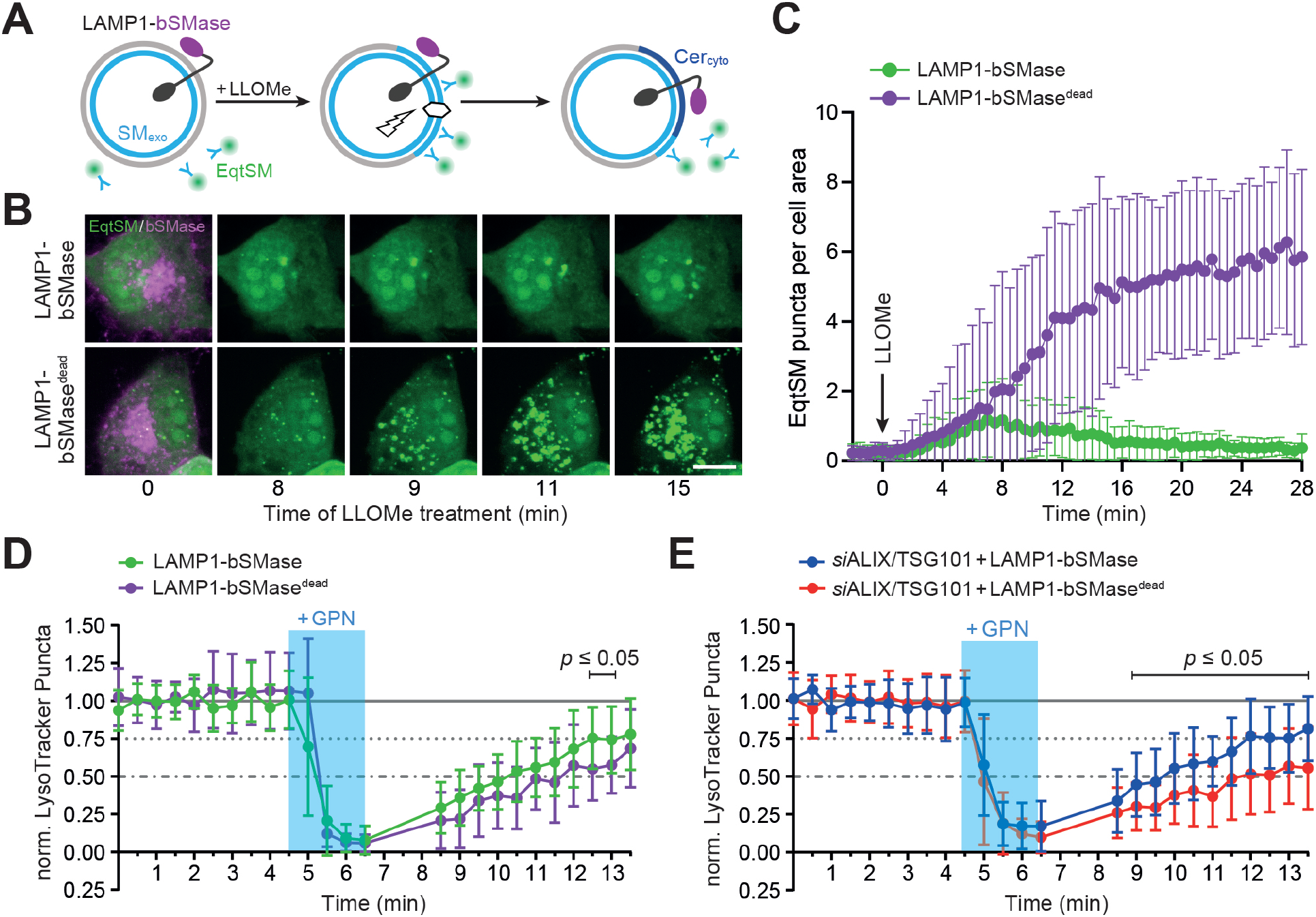
Hydrolysis of cytosolic SM promotes lysosomal repair in ESCRT-compromised cells. (**A**) Bacterial SMase (bSMase, *magenta*) was fused to the *C*-terminus of lysosomal membrane protein LAMP1, enabling an efficient metabolic turn-over of SM translocated to the cytosolic surface of LLOMe-damaged lysosomes. (**B**) Time-lapse images of HeLa cells co-expressing GFP/V5-tagged LAMP1-bSMase or LAMP1-bSMase^dead^ (*magenta*) and mKate-tagged EqtSM (*green*) treated with 1 mM LLOMe for the indicated time. Scale bar, 10 μm. (**C**) Time-course plotting EqtSM-positive puncta per 100 μm^2^ cell area in cells treated as in (B). Data are means ± SD from 14 cells per condition, *n* = 2. (**D**) Time-course plotting LysoTracker-positive puncta in HeLa cells co-expressing GFP/V5-tagged LAMP1-bSMase or LAMP1-bSMase^dead^ and mKate-tagged EqtSM during and after a 2 min-pulse of GPN, normalized to the initial number of puncta. Data are means ± SD from ≥ 16 cells per condition, *n* = 3. (**E**) Time-course plotting LysoTracker-positive puncta in HeLa cells pre-treated with siRNAs targeting GFP or ALIX/TSG101 (72 h) and expressing LAMP1-bSMase or LAMP1-bSMase^dead^ during and after a 2 min-pulse of GPN. Data are means ± SD from ≥ 23 cells per condition, *n* = 3. Statistical significance was determined by unpaired two-tailed t-test.

### Inhibition of neutral SMases disrupts lysosomal repair

As ectopic expression of a bacterial SMase targeted to the cytosolic surface of lysosomes enhanced lysosomal repair, we anticipated that inhibition of endogenous neutral SMases, which act on cytosolic SM pools, might perturb lysosomal repair. Indeed, addition of the generic neutral SMase inhibitor GW4869 significantly impaired the recovery of LysoTracker fluorescence in cells transiently exposed to GPN (Fig. 6A). The negative impact of GW4869 on lysosomal repair was even more pronounced in cells in which ESCRT function was compromised by pre-treatment with ALIX and TSG101-targeting siRNAs (Fig. 6B). Consistent with a critical role of neutral SMases in lysosomal repair, GW4869 significantly reduced the viability of both control and ALIX/TSG101-depleted cells in the presence of LLOMe (Fig. 6C, D). This raised the question which of the known neutral SMase isoforms participates in the restoration of damaged lysosomes. Four mammalian neutral SMases have been identified to date, namely nSMase-1 (SMPD2), nSMase-2 (SMPD3), nSMase-3 (SMPD4) and mitochondria-associated MA-nSMase (SMPD5)^27^. While expression of nSMase-3 is mainly restricted to skeletal muscle and heart^28^, nSMase-1 and −2 are ubiquitously expressed and promote SM hydrolysis on the cytosolic surface of the ER and plasma membrane, respectively^29,30^. Using the LysoTracking fluorescence recovery assay, we found that siRNA-mediated depletion of nSMase-1 had no impact on the repair of GPN-damaged lysosomes in ESCRT-compromised cells (Fig. 6E). In contrast, siRNA-mediated depletion of nSMase-2 significantly impaired the recovery of lysosomes injured by GPN (Fig. 6F; Fig. S8). This indicates that nSMase-2 is a critical component of a SM-dependent membrane repair pathway for the restoration of damaged lysosomes.

**Fig. 6.**
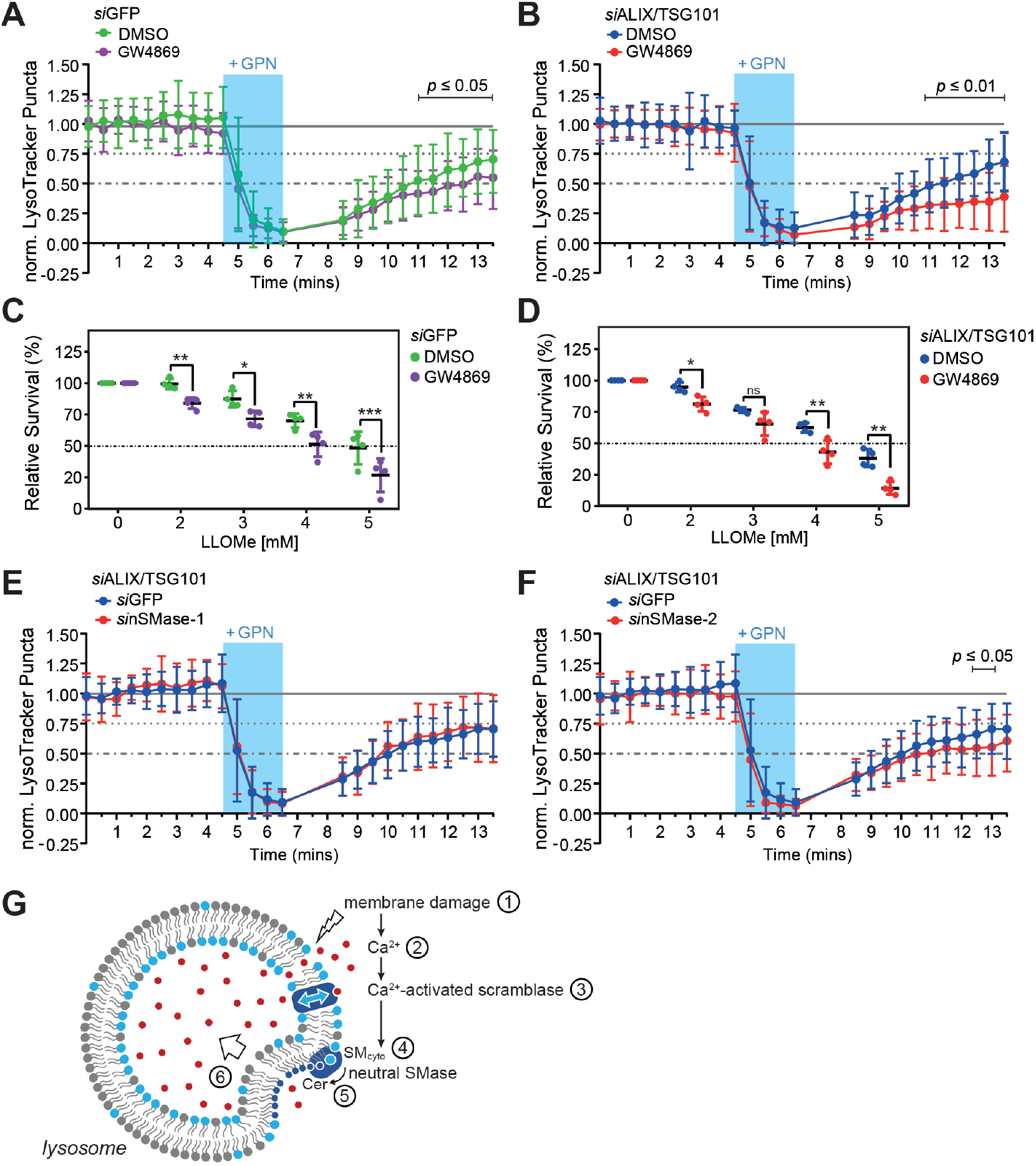
Inhibition of neutral SMases disrupts repair of damaged lysosomes. (**A**) Time-course plotting LysoTracker-positive puncta in HeLa cells pre-treated with siRNA targeting GFP (72 h) and 10 μM GW4869 or 0.5% (v/v) DMSO as vehicle control (30 min) during and after a 2 min-pulse of GPN. Data are means ± SD from ≥ 39 cells per condition, *n* = 3. (**B**) Time-course plotting LysoTracker-positive puncta in HeLa cells pre-treated with siRNAs targeting ALIX/TSG101 and 10 μM GW4869 or 0.5% (v/v) DMSO (30 min) during and after 2 min-pulse of GPN. Data are means ± SD from ≥ 44 cells per condition, *n* = 3. (**C**) Survival rate of HeLa cells pre-treated with siRNA targeting GFP (72 h) after 5 h exposure to LLOMe at the indicated concentration in the presence of 10 μM GW4869 or 0.5% (v/v) DMSO. Data are means ± SD, *n* = 3. (**D**) Survival rate of HeLa cells pre-treated with siRNA targeting ALIX/TSG101 (72 h) after 5 h exposure to LLOMe at the indicated concentration in the presence of 10 μM GW4869 or 0.5% (v/v) DMSO. Data are means ± SD, *n* = 3. (**E**) Time-course plotting LysoTracker-positive puncta in HeLa cells pre-treated with siRNAs targeting ALIX/TSG101 and GFP or nSMase-1 (72 h) during and after a 2 min-pulse of GPN. Data are means ± SD from ≥ 28 cells per condition, *n* = 3. (**F**) Time-course plotting LysoTracker-positive puncta in HeLa cells pre-treated with siRNAs targeting ALIX/TSG101 and GFP or nSMase-2 (72 h) during and after a 2 min-pulse of GPN. Data are means ± SD from ≥ 29 cells per condition, *n* = 3. Statistical significance was determined by paired (C,D) or unpaired two-tailed t-test (A,B,E,F). *p*-values are presented where significant. \**p* ≤ 0.05, \*\**p* ≤ 0.01, \*\*\**p* ≤ 0.001.

## DISCUSSION

The ESCRT machinery plays a well-established role in responding to and repairing damaged lysosomes. Here, we uncovered a complementary sphingolipid-operated lysosomal repair pathway that reverses potentially lethal membrane damage inflicted by lysosomotropic peptides and restores compartmental pH independently of ESCRT. The main features of this membrane repair pathway are depicted in Fig. 6G. We envision that minor perturbations in the integrity of the lysosomal membrane cause calcium ions to leak from the organelle’s lumen into the cytosol. A local rise in cytosolic calcium triggers calcium-activated scramblases near the injury site, resulting in a rapid exposure of SM on the cytosolic surface of the damaged organelle. The expanding pool of SM in the cytosolic leaflet is then turned-over by a neutral SMase, presumably nSMase-2, which cleaves the bulky phosphorylcholine head group of SM to generate ceramide, a lipid with a much smaller and less-hydrated head group. Ceramide has a cone-shaped structure and occupies a smaller membrane area than SM. Ceramides released by SM turnover self-assemble into microdomains that possess a negative spontaneous curvature^31^, causing a local condensation of the cytosolic leaflet. This, in turn, would promote an inverse budding of the bilayer away from the cytosol, akin to the ESCRT-mediated formation of intraluminal vesicles^32^. Collectively, our data indicate that a calcium-induced SM scrambling and turnover drives an ESCRT-independent mechanism to clear minor lesions from the lysosome-limiting membrane and prevent lysosomal damage-induced cell death. In line with our findings, ceramide-based membrane invaginations were demonstrated in SM-containing giant liposomes exposed to external SMases^33,34^ and have been implicated in the biogenesis of proteolipid-containing exosomes inside multivesicular endosomes, a process occurring independently of ESCRT and with nSMase-2 playing a crucial role^35,36^.

Lysosomal acid SMase (aSMase) previously emerged as a key player in the repair of plasma membrane damage caused by pore-forming toxins. Thus, aSMase has been shown to promote plasma membrane invagination and endocytosis in SLO-permeabilized cells in response to Ca^2+^-triggered exocytosis of lysosomes^37^. Here, removal of SLO-damaged plasma membrane areas relies on ceramide microdomain formation in the exoplasmic leaflet through aSMase-mediated hydrolysis of SM, which is normally concentrated there. While this process is mediated by a classical budding of the bilayer toward the cytosol, there is also evidence for alternative plasma membrane repair pathways in which lesions are removed by a reverse budding and shedding of extracellular vesicles. For instance, real-time imaging and correlative scanning electron microscopy of cells wounded by a laser provided evidence for ESCRT-mediated extracellular shedding of the damaged plasma membrane area^7^. Moreover, recent work revealed that TMEM16F promotes plasma membrane repair in cells exposed to pore-forming toxins by facilitating the release of extracellular vesicles to eliminate the toxin from the membrane^38^. This raised the idea that PS exposure catalyzed by TMEM16F helps protect cells from external attacks and injuries by constituting a “repair me” signal. However, our present findings raise an alternative scenario in which an injury-induced scrambling of SM mediated by TMEM16F serves to fuel a sphingolipid-based membrane restoration pathway analogous to the one that operates in lysosomes. Thus, the striking SM asymmetry that marks late secretory and endolysosomal organelles may actually reflect a vital role of SM and neutral SMases in safeguarding the functional integrity of these organelles.

## ONLINE METHODS

### Chemical reagents

Chemical reagents were used at the following concentrations, unless indicated otherwise: 1 mM LLOMe (Bachem; 4000725); 200 μM GPN (Abcam; ab145914); 1500 U/ml SLO (Sigma-Aldrich; S5265); 250 μM digitonin (Sigma-Aldrich; D141); 5 μM ionomycin (Sigma-Aldrich; I0634); 100 μM BAPTA-AM (Cayman Chemical; 15551); 75 nM LysoTracker™ Red DND-99 (Thermo Fisher Scientific; L7528); 1 μg/ml doxycycline (Sigma-Aldrich;); and 10 μM GW4869 (Sigma-Aldrich; D1692). GW4869 was stored at −80°C as a 2 mM stock suspension in dimethyl sulfoxide (DMSO). Just before use, the suspension was solubilized by addition of 5% methane sulfonic acid as described in (*39*).

### Antibodies

Antibodies used were: rabbit polyclonal anti-TMEM16F (Sigma-Aldrich; HPA038958; IB 1:1000); mouse monoclonal anti-SMS2 (Santa Cruz; sc-293384; IB 1:1000); rabbit polyclonal anti-CHMP4B (Proteintech; 13683-1-AP; IF 1:300); mouse monoclonal anti-ALIX (Biolegend; 634501; IB 1:1000); mouse monoclonal anti-Actin (Sigma-Aldrich; A1978; IF 1:1200; IB 1:10,000); rabbit monoclonal anti-Na/K-ATPase (Abcam; ab-76020; IF 1:600); mouse monoclonal anti-TSG101 (Santa Cruz; sc-7964; IB 1:1000); mouse monoclonal anti-V5 (Invitrogen; r96025; IF 1:400; IB 1:1000); HRP-conjugated goat anti-mouse IgG (Thermo Fisher Scientific; 31430; IB 1:5000); Cy™2-conjugated donkey anti-mouse IgG (Jackson ImmunoResearch Laboratories; 715-225-150; IF 1:400); and Cyanine Cy™3-conjugated donkey anti-rabbit IgG (Jackson ImmunoResearch Laboratories; 715-165-152; IF 1:400).

### DNA constructs

Expression constructs encoding cytoplasmic GFP-tagged EqtSM and EqtSol were created by PCR amplification of DNA encoding residues 22-227 of Eqt-SM or Eqt-Sol^*40*^ followed by cloning into EcoRI and BamHI sites of pN1-oxGFP^*17*^. Expression constructs encoding cytoplasmic mKate-tagged EqtSM and EqtSol were created by PCR amplification followed by cloning into NheI and AgeI sites of mKate-LifeAct-7 (Addgene #54697), thereby replacing the LifeAct ORF. Expression constructs encoding mCherry-tagged human Galactin3 (pLX304-mCherry-hGalectin3), GFP-tagged human LAMP1 (pCMV-hLAMP1-GFP) and mCherry-tagged human LAMP1 (pCMV-hLAMP1-mCherry) were kindly provided by Michael Hensel (University of Osnabrück, DE) and have been described in (*41,42*). Expression constructs encoding bacterial SMase fused to GFP-tagged LAMP1 were created by PCR amplification of DNA encoding residues 28-333 of *Bacillus cereus* SMase using pEF6-bSMase-V5-His and pEF6-bSMase^D322A/H323A^-V5-His^43^ as templates, followed by cloning into BamHI and AgeI sites of pCMV-hLAMP1-GFP to yield LAMP1-bSMase-V5-GFP and LAMP1-bSMase^dead^-V5-GFP. Expression construct pEF6-nSMase2-V5 encoding V5-tagged human neutral SMase2 has been described in (*44*). A doxycycline-inducible expression construct encoding human SMS1 with a *N*-terminal FLAG tag (MDYKDDDDK) was created by PCR amplification using pcDNA1.3-SMS1-V5^45^ as a template, followed by cloning into BamHI and NotI sites of pENTR™11 (Invitrogen; A10467). The insert was next transferred into lentiviral expression vector *pInducer20* (Addgene #44012) using Gateway cloning, according to the manufacturer’s instructions.

### Cell culture and siRNA treatment

Human cervical carcinoma HeLa (ATCC CCL-2) and human embryonic kidney HEK293T cells (ATCC CRL-3216) were cultured in Dulbecco’s modified Eagle’s medium (DMEM) supplemented with 10% FBS (Pan Biotech; P40-47500). Murine RAW264.7 macrophages (ATCC TIB-71) were cultured in RPMI supplemented with 10% FBS. A HeLa cell-line stably expressing CHMP4B-eGFP was kindly provided by Anthony Hyman (Max Planck Institute for Molecular Cell Biology and Genetics, Dresden, DE) and has been described in^46^. DNA transfections were performed using Lipofectamine 3000 (Thermo Fisher). Treatment with siRNAs (Qiagen) were carried out using Oligofectamine reagent (Invitrogen) according to the manufacturer’s instructions. siRNA target sequences were: GFP, 5’-GCACCATCTTCTTCAAGGACG-3’; ALIX, 5’-CCUGGAUAAUGAUGAAGGA-3’; TSG101, 5’-CCUCCAGUCUUCUCUCGUC-3’; nSMase1, 5’-CAGCAGAGAGGUCGCCGUU-3’; nSMase2, 5’-CAAGCGAGCAGCCACCAAA-3’.

### RT-qPCR

To verify siRNA-mediated knock-down of gene expression, RNA was extracted from siRNA-treated cells using TRIzol reagent (Thermo Fisher Scientific). One μg of RNA was used to synthesize cDNA with the Superscript III Reverse Transcriptase Kit (Thermo Fisher Scientific; 18080051) according to the manufacturer’s instructions. Quantitative PCR reactions were performed on a C1000 Thermal Cycler with a CFX96 Real-Time System (Bio-Rad) using Maxima™ SYBR™ Green/ROX 2x qPCR Master Mix (Thermo Fisher Scientific; K0221). Each reaction contained 400 ng of cDNA and 0.3 μM each of sense and anti-sense primers in a total volume of 10 ul. Initial denaturation was at 95°C for 10 min. Cycles (n = 40) consisted of 10 s denaturation at 95°C, 30 s annealing at 57°C and 30 s extension at 72°C. Analysis of a single PCR product was confirmed by melt-curve analysis. All reactions were performed in triplicate. Expression data were normalized using actin as a reference. Ct values were converted to mean normalized expression using the 2^-ΔΔCt^ method (Livak and Schmittgen, Methods, 2001). Primers used were: nSMase1-sense, 5’-GGTGCTCAACGCCTATGTG-3’; nSMase1-antisense, 5’-CGTCTGCCTTCTTGGATGTG-3’; nSMase2-sense, 5’-CAACAAGTGTAACGACGATGCC-3’; nSMase2-antisense, 5’-CGATTCTTTGGTCCTGAGGTGT-3’; actin-sense, 5’-ATTGGCAATGAGCGGTTCC-3’; actin-antisense, 5’-GGTAGTTTCGTGGATGCCACA-3’.

### Generation of TMEM16F-KO cells

To knock out TMEM16F in HeLa cells, a mix of CRISPR/Cas9 constructs encoding three different TMEM16F-specific gRNAs and a GFP marker was obtained from Santa Cruz (sc-402736). The TMEM16F specific gRNA sequences were: A/sense, 5’-CAGCCTTTGGTACACTCAAC-3’; B/sense, 5’-GAATCTAACCTTATCTGTCA-3’; C/sense, 5’- AATAGTACTCACAAACTCCG-3’. At 24 h post-transfection, GFP-positive single cells were sorted into 96 well plates using a SH800 Cell Sorter (Sony Biotechnology), expanded and analyzed for TMEM16F expression by immunoblot analysis. In addition, loss of TMEM16F function was verified by analyzing ionomycin-treated cells for surface exposure of phosphatidylserine using Annexin V staining. To this end, wild type and TMEM16F-KO HeLa cells were detached using trypsin, taken up in DMEM containing 10% FBS, washed in PBS and resuspended in Annexin V Binding Buffer (Biolegend, no. 422201) and then incubated in the presence or absence of 15 μM ionomycin for 10 min at 37°C in 5% CO_2_. Next, APC-Annexin V (Biolegend, no. 640920; 5 μl in 100 μl Binding Buffer) and propidium iodide (5 μg/ml; Sigma Aldrich, no. P4170) were added and cells were incubated for 10 min at RT. After addition of 400 μl Annexin V Binding Buffer, cells were cooled on ice and then subjected to flow cytometry using a SH800 Cell Sorter (Sony Biotechnology).

### Generation of SMS-KO cells

To generate SMS1 and SMS2 double-KO (SMS-KO) HeLa cells, a mix of CRISPR/Cas9 constructs encoding three different gRNA per gene and the corresponding HDR plasmids were obtained from Santa Cruz (SMS1, sc-403382; SMS2, sc-405416). SMS1-specific gRNA sequences were: A/sense, 5’- TG ATACCACCAGAGTCGCCG-3’; B/sense, 5’- TTGTACCTCGATCTTACCAT-3’; C/sense, 5’- TAAGTGTTAGCATGACCGTG-3’. SMS2-specific gRNA sequences were: A/sense, 5’-TAACCGTGTGACCGCTGAAG-3’; B/sense, 5’-GGTCTTGCATAAGTGTTCGT-3’; C/sense, 5’- GTTACTACTCTACCTGTGCC-3’. Cells transfected with the SMS2-KO constructs were grown in medium containing 2 μg/ml puromycin at 48 h post-transfection (Sigma-Alderich; P8833). After 1-2 weeks, single drug-resistant colonies were picked, expanded and analysed for SMS2 expression by immunoblot analysis using a mouse monoclonal anti-SMS2 antibody. A SMS1/2 double-KO cell line (clone #7) was created by transfecting SMS2-KO clone #25 with SMS1-KO constructs as above, after ejection of the puromycin selectable marker using Cre vector (Santa Cruz; sc-418923) according to the manufacturer’s instructions. Loss of SMS1 was confirmed by metabolic labeling of double-KO candidates with 4 μM of the clickable sphingosine (2*S*, 3*R*, 4*E*)-2-Amino-octadec-4-en-17-yne-1,3-diol (clickSph) in Opti-MEM (Fisher Scientific Scientific; 11520386) for 24 h. The synthesis of clickSph will be described elsewhere (S. Korneev and J. Holthuis, unpublished data). Next, cells were washed in PBS, harvested, and subjected to Bligh and Dyer lipid extraction (Bligh and Dyer, 1959). Dried lipid films were click reacted with the fluorogenic dye 3-azido-7-hydroxycoumarin (Jena Bioscience; CLK-FA047) by addition of 64.5 μl of a freshly prepared click reaction mix containing 0.45 mM 3-azido-7-hydroxycoumarin and 1.4 mM Cu(I)tetra(acetonitrile) tetrafluoroborate in CH_3_CN:EtOH (3:7, v:v) for 2.5 h at 45°C without shaking. The reaction was quenched by addition of 150 μl methanol, dried down in a Speed-Vac, dissolved in CHCl_3_:methanol (2:1, v:v), and applied at 120 nl/s on a NANO-ADAMANT HP-TLC plate (Macherey-Nagel, Germany) with a CAMAG Linomat 5 TLC sampler (CAMAG; Switzerland). The TLC plate was developed in CHCl_3_:MeOH:H_2_O:AcOH (65:25:4:1, v:v:v:v) using a CAMAG ADC2 automatic TLC developer (CAMAG; Switzerland). Fluorescent lipids were analyzed using a ChemiDoc XRS+ with UV-transillumination and Image Lab Software (BioRad, USA).

### Lentiviral transduction

HeLa SMS-KO cells with a stably integrated doxycycline-inducible SMS1 expression construct were created by lentiviral transduction. To this end, HEK293T cells were co-transfected with *pInducer20*-FLAG-SMS1 and the packaging vectors psPAX2 (Addgene #12260) and pMD2.G (Addgene #12259). Culture medium was changed 6 h post-transfection. After 48 h, the lentivirus-containing medium was harvested, passed through a 0.45 μm filter, mixed 1:1 (v/v) with DMEM containing 8 μg/ml polybrene (Sigma-Aldrich; TR-1003) and used to infect HeLa SMS-KO cells. At 24 h post-infection, the medium was replaced with DMEM containing 1 mg/ml G418 (Sigma-Aldrich; G8168) and selective medium was changed daily. After 3-5 days, positively transduced cells were selected and analyzed for doxycycline-dependent expression of FLAG-SMS1 by immunoblot analysis, immunofluorescence microscopy, and metabolic labeling with clickSph as described above.

### SMase activity assay

HeLa cells were seeded in a 6-well plate at 150.000 cells per well in DMEM supplemented with 10% FBS. After 24 h, cells were transfected with nSMase2-V5, LAMP1-bSMase-V5-GFP or LAMP1-bSMase^dead^-V5-GFP and grown for 24 h. Next, cells were harvested in ice-cold lysis buffer (25 mM Tris pH 7.4, 0.1 mM PMSF, 1x protease inhibitor cocktail), subjected to sonication (Branson Ultrasonic Sonifier) and centrifuged at 500 *g* for 10 min at 4°C to obtain a post-nuclear fraction. Aliquots equivalent to 20 μg of total protein were included in a 100 μl reaction mixture containing 50 mM Tris pH 7.4, 10 mM MgCl_2_, 0.2% Triton X-100, 10 mM DTT, 50 μM phosphatidylserine (Sigma-Aldrich; P7769) and 50 μM C_6_-NBD-SM (Biotium; 60031). Reactions were incubated at 37°C for 2 h, terminated by addition of MeOH/CHCl_3_, subjected to a Bligh and Dyer lipid extraction and then analyzed by TLC as described above.

### Cytotoxicity assay

Cells treated with siRNAs were seeded in a 96-well plate (Greiner Bio-One; 655101) at 10.000 cells per well in DMEM supplemented with 10% FBS at 24 h after starting the treatment. After 24 h, the medium was replaced with Opti-MEM, and 24 h later LLOMe was added at the indicated concentration. After 3.5 h, PrestoBlue HS (Thermo Fisher Scientific; P50200) was added directly to the well to a final concentration of 10% (v/v) and incubated for 1.5 h at 37°C. Next, absorbance at 570 nm was measured with 600 nm as reference wavelength using an Infinite 200 Pro M-Plex plate reader (Tecan Lifesciences). To calculate relative percentage of survival, the measured value for each well (x) was subtracted by the minimum measured value (min) and divided by the subtrahend of the average measured value of untreated cells (untreated) and the minimum measured value (min); ((x-min)-(untreated-min)). To analyse the impact of GW4869 on LLOMe sensitivity, cells were seeded in a 96-well plate as above and grown for 48 h. Next, cells were treated with 10 μM GW4869 or 0.5% DMSO (vehicle control) for 30 min before LLOMe was added at the indicated concentration. Cell viability was assessed as described above.

### Time-lapse recording of laser wounding

Laser wounding and time-lapse acquisition were performed using an Olympus model FV3000 laser scanning microscope (Olympus Europa SE & CO. KG) optically coupled to a fs laser system that comprises a regeneratively amplified fs laser (Pharos-HE-20; Light Conversion Inc.) and an optical parametric amplifier (OPA, Orpheus-Twins F; Light Conversion Inc.). The fs-pulses are adjusted collinear to the optical path of the continuous laser integrated in the microscope, enabling the simultaneous use of the continuous and pulsed laser as pumping source. Prior to the coupling into the microscope, neutral density filters can be inserted to tailor the average power and so the energy per pulse. The filter combination for simultaneous IR and blue emission was BP 488/730-1200. The wavelength of the pulsed laser was set at 900 nm, the repetition rate was 10 kHz, and the pulse duration was 180 fs, with a total exposure time of 3 sec. The average power at the sample position was 900 μW, which implies that an energy per pulse of 90 nJ was achieved. For time-lapse recording of laser wounding, an UPLSAPO 60x water immersed NA 1.2 objective with a custom infrared (IR) coating was used. Based on the diffraction limited spot created by the objective, the photodamaged area was estimated to be approximately 500 nm. For laser wounding experiments, HeLa cells were seeded on 24 mm glass coverslips in a 6-well plate at a density of 150.000 cells per well in DMEM containing 10% FBS. After 24 h, the medium was replaced with Opti-MEM and cells were transfected with EqtSM-GFP or EqtSol-GFP. After 24 h, Opti-MEM was replaced with Imaging Medium (IM; 30 mM HEPES, 140 mM NaCl, 2.5 mM KCl, 1 mM MgCl_2_, 1.8 mM CaCl_2_, 10 mM D-glucose, pH7.4.) and cells were transferred to the microscope. Time-lapse images were acquired every 5 sec before and after laser wounding (5 z-sections, 1 μm apart).

### Time-lapse recordings of cells exposed to organelle-damaging agents

Time-lapse recordings of cells exposed to organelle-damaging drugs or pathogens were performed using a Zeiss Cell Observer Spinning Disc Confocal Microscope equipped with a TempModule S1 temperature control unit, a Yokogawa Spinning Disc CSU-X1a 5000 Unit, a Evolve EMCDD camera (Photonics, Tucson), a motorized xyz-stage PZ-2000 XYZ (Applied Scientific Instrumentation) and an Alpha Plan-Apochromat x 63 (NA 1.46) oil immersion objective. The following filter combinations were used: blue emission with BP 445/50, green emission with BP 525/50, orange emission BP 605/70. All images were acquired using Zeiss Zen 2012 acquisition software. At 48 h before imaging, cells were seeded into a μ-Slide 8 well glass bottom chamber (Ibidi; 80827) at a density of 20.000 cells per well in DMEM supplemented with 10% FBS. After 24 h, the medium was replaced with Opti-MEM and cells were transfected with expression constructs encoding fluorescently-tagged proteins. After another 24 h, Opti-MEM was replaced with IM containing 30 mM HEPES, 140 mM NaCl, 2.5 mM KCl, 1 mM MgCl_2_, 1.8 mM CaCl_2_ and 10 mM D-glucose, pH7.4. Next, cells were immediately transferred to the stage-top incubator preheated to 37°C. The slide was allowed to equilibrate for 10 min before initiation of image acquisition. For experiments under Ca^2+^-depleted conditions, CaCl_2_-free IM was used, which was supplemented with either 2 mM EGTA or 100 μM BAPTA-AM. Time-lapse images were acquired every 10-30 sec (6 z-sections, 1 μm apart). After 2 min, organelle-damaging agents were added directly to the well without pausing image acquisition.

### *M. marinum* infection

RAW264.7 cells were seeded into a SensoPlate™ 96-Well Glass-Bottom Plate (Greiner Bio-One; M4187) at a density of 10.000 cells per well in RPMI supplemented with 10% FBS. After 24 h, cells were transfected with EqtSM-GFP or EqtSol-GFP. At 24 h post-transfection, cells were infected with *M. marinum* wildtype or ΔRD1 mutant strains constitutively expressing mCherry at an MOI of 10. Strains were kindly provided by Caroline Barisch (University of Osnabrück) and have been described in^47^. The 96-well plate was centrifuged at 1250 *g* for 30 sec and then incubated for 2 h at 37°C. Next, cells were washed with PBS and fixed with 4% (w/v) paraformaldehyde (PFA) in PBS for 15 min at RT. For time-lapse imaging, cells were grown in phenol red-free RPMI medium supplemented with 30 mM HEPES and infected with the above *M. marinum* strains at an MOI of 25. After centrifugation of the 96-well plate at 1250 g for 30 secs, cells were imaged using the Zeiss Cell Observer SD microscope set-up with images captured at 1 min time intervals (5 z-sections, 1μm apart).

### Immunostaining of fixed cells

For treatment with digitonin, cells were seeded onto 12 mm glass coverslips at a density of 40.000 cells per coverslip in DMEM supplemented with 10% FBS. After 24 h, the medium was replaced with Opti-MEM and cells were transfected with EqtSM-GFP. After 48 h, cells were treated with 250 μM digitonin for 1 min, washed twice in Opti-MEM, incubated in Opti-MEM for 3 min at 37°C and then fixed with 4% (w/v) PFA in PBS for 15 min at RT. After quenching in 50 mM ammonium chloride, cells were permeabilized using PBS containing 0.1% (w/v) saponin and 0.2% (w/v) BSA and immunostained for Na/K-ATPase and counterstained with DAPI. For monitoring recruitment of endogenous CHMP4B to LLOMe-damaged lysosomes, cells were seeded onto 12 mm glass coverslips at a density of 40.000 cells per coverslip in DMEM supplemented with 10% FBS. After 24 h, the medium was replaced with Opti-MEM. At 48 h post-seeding, cells were incubated with Opti-MEM containing 1 mM LLOMe for the indicated time period, washed with PBS, and then fixed with MeOH at −20°C for 15 min. After fixation, cells were washed three times with PBS and permeabilized in PBS containing 0.3% (v/v) Triton-X100 and 1% (w/v) BSA for 15 min. Cells were immunostained for CHMP4B and actin, and counterstained with DAPI.

### Time-lapse recordings of LysoTracker-labeled cells

At 72 h before imaging, cells were treated with siRNAs as indicated. At 48 h before imaging, cells were seeded in a μ-Slide 8 well glass bottom chamber (Ibidi; 80827) at a density of 20.000 cells per well in DMEM supplemented with 10% FBS. After 24 h, the medium was replaced with Opti-MEM and cells were transfected with expression constructs encoding fluorescently-tagged proteins as indicated. At 24 h post-transfection, Opti-MEM was replaced by IM containing 75 nM LysoTracker (LT) and the cells were immediately transferred to a stage-top incubator preheated to 37°C. The slide was allowed to equilibrate for 10 min before initiation of image acquisition with the Zeiss Cell Observer SD microscope. Time-lapse images were acquired every 30 sec (6 z-sections, 1 μm apart). After mins of image acquisition, GPN was directly added into the well to a final concentration of 200 μM without pausing the acquisition. After 2 min of GPN exposure, acquisition was paused for 2 min to aspirate the GPN-containing medium, wash the cells once with LT-containing IM and add fresh LT-containing IM before acquisition was resumed. To analyse the impact of GW4869 on the recovery of LysoTracker fluorescence after transient GPN exposure, cells were seeded in a μ-Slide 8 well glass bottom chamber, subjected to medium changes as above and incubated for 48 h. Next, the cells were treated with 10 μM GW4869 or 0.5% DMSO (vehicle control) for 10 min in IM without LT and subsequently for 10 min in IM containing 75 nM LT. The GW4869 and DMSO concentrations were kept constant during and after the 2-min GPN exposure and images were acquired every 30 sec as described above.

### Image analysis

All image analyses were performed using Image J macros on the original, unmodified data. Only cells that maintained a healthy morphology were included into the analysis. To quantify the number of EqtSM, Gal3 or CHMP4B-positive puncta during time-lapse imaging, the background was subtracted to remove noise and a manual threshold was set to exclusively include puncta above the signal of the cytosolic probe in untreated cells. Puncta with close proximity were separated using the watershed function. Next, for each time point, all puncta with pre-determined characteristics were counted automatically (size 0.2-5 μm^2^, circularity 0.5-1). For cells co-expressing CHMP4B-eGFP and EqtSM-mKate, the nuclear area was excluded from the analysis. For normalization, the number of puncta for each time point was divided by the total measured area to account for size difference of cells and then multiplied by 100 to obtain the number of puncta per 100 μm^2^ cell area. For normalization relative to the maximal value, the maximum number of puncta for each cell was determined and each time point was divided by the maximum value. To quantify the intensity of EqtSM-positive puncta upon laser damage, the images were first corrected for bleaching. Next, an ROI at the site of damage was selected and after a fixed threshold was implemented with the “Minimum” setting, the relative intensity in the ROI was measured for each time point. For quantifying LT-positive puncta, the background was subtracted to remove noise and an automatic threshold was set for t = 0 min. Puncta with close proximity were separated using the watershed function. Next, for each time point, all puncta with pre-determined characteristics were counted automatically (size 0.2-5 μm^2^, circularity 0.5-1). For quantification, the average of the first five time points (0 - 2min) was calculated and every time point was divided by the average. To quantify the LT accumulation efficiency, the background was subtracted to remove noise and an automatic threshold was set for t = 10 min. Puncta with close proximity were separated and for each time point, puncta with pre-determined characteristics were counted automatically (size 0.2-5μm^2^, circularity 0.5-1). For quantification, the average for the time points from t = 8 to t = 10 min was calculated and every time point was divided by the average. To quantify the intensity of the CHMP4B immunostaining in LLOMe-treated cells, the cell outline marked by actin immunostaining was used to measure the cell area. For the CHMP4B immunostaining, a pre-determined, fixed threshold was applied and the intensity above the threshold was measured. The intensity above threshold was divided by the cell area. The average value for 10 min LLOMe treatment was set to 1 and all data points were divided by the average value. Each individual measurement was plotted in a violin plot.

### Image J Macros

#### Quantification LysoTracker puncta

1. run(“Subtract…”, “value=10 stack”);
2. setAutoThreshold(“Default dark no-reset”);
3. setOption(“BlackBackground”, false);
4. run(“Convert to Mask”, “method=Default background=Dark”);
5. run(“Convert to Mask”, “method=Default background=Light”);
6. run(“Watershed”, “stack”);
7. run(“Analyze Particles…”, “size=0.2-5 circularity=0.5-1.00 show=Outlines display exclude summarize stack”);
8. run(“Next Slice [>]”)
9. // repeat step 7 and 8 until all time frames have been quantified

#### CHMP4B recruitment / Pre-processing

1. imageTitle=getTitle();
2. run(“Split Channels”);
3. selectWindow(“C1-”+imageTitle);
4. rename(“DAPI”);
5. selectWindow(“C2-”+imageTitle);
6. rename(“Actin”);
7. selectWindow(“C3-”+imageTitle);
8. rename(“CHMP4B”);
9. run(“Merge Channels…”, “c2=[Actin] c1=[CHMP4B] c3=[DAPI] create”);
10. run(“Z Project…”, “projection=[Max Intensity]”);
11. run(“Split Channels”);
12. selectWindow(“C1-MAX_Composite”);
13. setMinAndMax(200, 2000);
14. run(“Subtract…”, “value=10”);
15. selectWindow(“C2-MAX_Composite”);
16. setMinAndMax(450, 2200);
17. run(“Subtract…”, “value=50”);
18. selectWindow(“C3-MAX_Composite”);
19. setMinAndMax(400, 700);
20. run(“Merge Channels…”, “c1=[C1-MAX_Composite] c2=[C2-MAX_Composite] c3=[C3-MAX_Composite] create”);
21. selectWindow(“Composite”);
22. close();

#### CHMP4B recruitment / Quantification

1. //setTool(“freehand”);
2. run(“Measure”);
3. run(“Duplicate…”, “duplicate”);
4. run(“Split Channels”);
5. close();
6. selectWindow(“C1-MAX_Composite-1”)
7. setAutoThreshold(“Default dark no-reset”);
8. //run(“Threshold…”);
9. setThreshold(1400, 65535);
10. run(“Convert to Mask”);
11. run(“Measure”);
12. selectWindow(“C2-MAX_Composite-1”);
13. close();

#### Quantification EqtSM-GFP and mCherry-Galectin3 puncta

1. //set Threshold manually
2. setOption(“BlackBackground”, false);
3. run(“Convert to Mask”, “method=Default background=Dark”);
4. run(“Fill Holes”,“stack”);
5. run(“Watershed”, “stack”);
6. run(“Analyze Particles…”, “size=0.2-5 circularity=0.50-1.00 show=Outlines display exclude summarize stack”);
7. run(“Next Slice [>]”)
8. //Repeat step 6 and 7 until all time points are analyzed

#### Quantification CHMP4B-GFP and EqtSM-mKate recruitment

1. //perform bleach correction with “exponential fit”
2. //select region that excludes nucleus
3. setOption(“BlackBackground”, false);
4. run(“Convert to Mask”, “method=Default background=Dark”);
5. run(“Fill Holes”,“stack”);
6. run(“Analyze Particles…”, “size=0.2-5 circularity=0.0-1.00 show=Outlines display exclude summarize stack”);
7. run(“Next Slice [>]”)
8. //Repeat step 6 and 7 until all time points are analyzed

## Acknowledgments

We gratefully acknowledge Caroline Barisch, Michael Hensel, Anthony Hyman, and Yussuf Hannun for providing DNA constructs, cell lines and bacterial strains, and Rainer Kurre for technical support in live cell microscopy.

## Author contributions

P. N. and J. C. M. H. designed the research and wrote the manuscript; P. N. performed the experiments and analyzed the results; T. S. and A. H. generated and analyzed the SMS-KO, TMEM16F-KO, and *pInd-*SMS1 cell lines; L. V. and M. I. assisted with 2-photon laser damage; Y. D., Y. K. and C. G. B. designed and characterized the equinatoxin probes; C. J. C. provided intellectual expertise and helped to interpret experimental results; All authors discussed results and commented on the manuscript.

## Data and materials availability

All data is available in the main text or Supplementary Materials.

