## Supplemental Information for "Ca^2+^-activated sphingomyelin scrambling and turnover mediate ESCRT-independent lysosomal repair"

**This PDF file includes:**

Figs. S1 to S8

Captions for Movies S1 to S7

**Other Supplementary Materials for this manuscript include:**

Movies S1 to S7

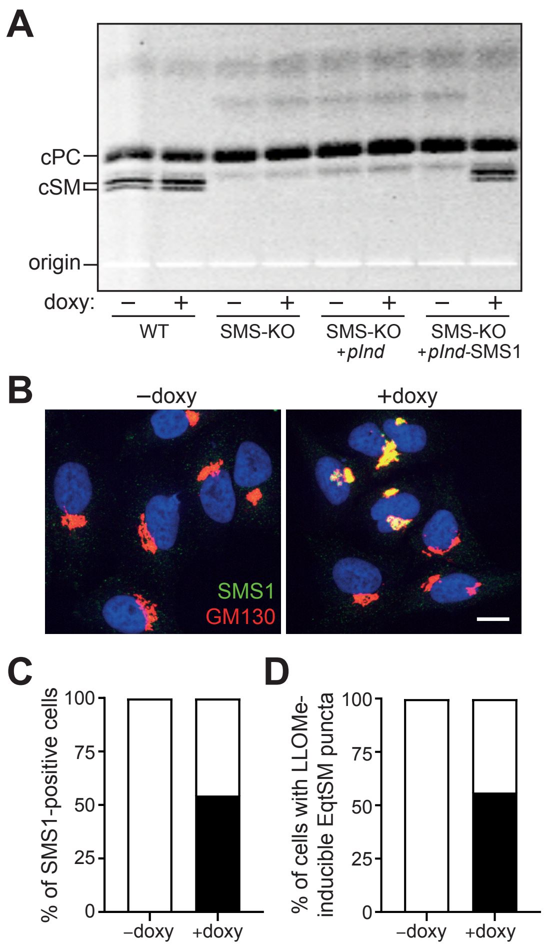


**Fig. S1. Characterization of SMS-KO cells transduced with doxycycline-inducible SMS1.**

(**A**) Wildtype (WT), SMS-KO or SMS-KO HeLa cells transduced with empty vector (*pInd*) or Flag-tagged SMS1 under control of a doxycycline-inducible promotor (*pInd*-SMS1) were cultured in the absence or presence of 1 µg doxycycline for 32 h. Next, cells were metabolically labeled with a clickable sphingosine analogue for 16 h in the absence (-) or presence (+) of doxycycline. Total lipids were extracted, click-reacted with 3-azido-7-hydroxycoumarin, separated by TLC, and analyzed by fluorescence detection. cPC, coumarin-labeled phosphatidylcholine; cSM, coumarin-labeled sphingomyelin. (**B**) HeLa SMS-KO cells transduced with *pInd*-SMS1 were cultured in the absence or presence of 1 mM doxycycline for 48 h, fixed, stained with antibodies against the Flag-tag (*green*) or the Golgi marker GM130 (*red*) and DAPI (*blue*), and then visualized by confocal microscopy. Scale bar, 10 µm. (**C**) Percentage of SMS-KO cells transduced with *pInd*-SMS1 and treated as in (B) displaying anti-Flag immunostaining. A minimum of 100 cells were analyzed per condition. (**D**) Percentage of SMS-KO cells transduced with *pInd*-SMS1 and expressing GFP-tagged EqtSM displaying GFP-positive puncta in response to GPN (200 mM, 2 min) after 48 h pre-incubation in the presence of absence of 1 µg/ml doxycycline. A minimum of 24 cells were analyzed per condition in three independent experiments.

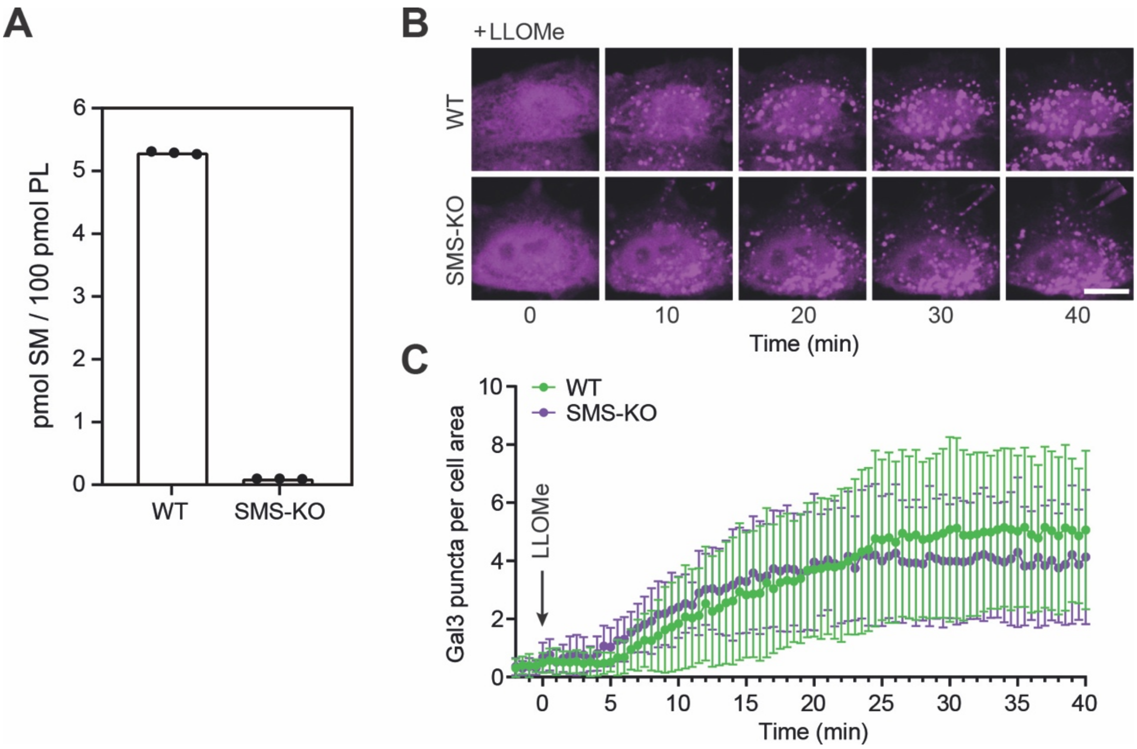


**Fig. S2. SM is dispensable for Gal3 recruitment to LLOMe-damaged lysosomes.**

(**A**) SM levels in Bligh and Dyer lipid extracts of wildtype (WT) or SMS-KO HeLa cells were determined by LC-MS/MS as in (*48*) and expressed in pmol per 100 pmol of total phospholipid analyzed. Data are means ± SD, $n$ = 3. (**B**) Time-lapse fluorescence micrographs of wildtype (WT) or SMS-KO HeLa cells expressing mCherry-tagged galectin-3 (Gal3) and treated with 1 mM LLOMe for the indicated time. (**C**) Time-course plotting Gal3-positive puncta per 100 µm$^2$ cell area in cells treated as in (A). Data are means ± SD from ≥ 8 cells per condition. Scale bar, 10 µm.

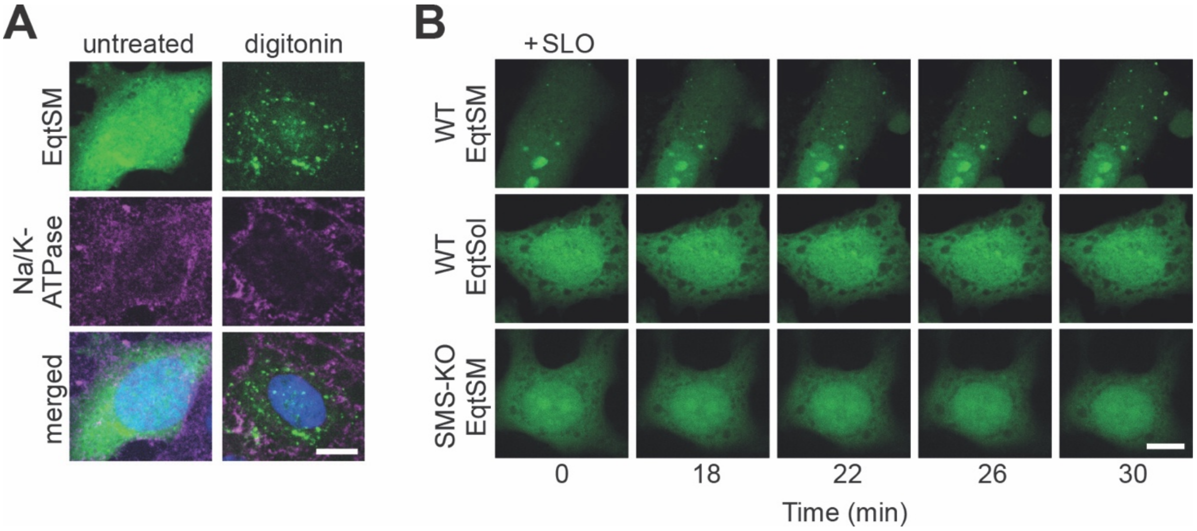


**Fig. S3. EqtSM is recruited to the plasma membrane damaged by pore-forming agents.**

(**A**) HeLa cells expressing GFP-tagged EqtSM were incubated in the absence or presence of 250 µM digitonin for 1 min. Next, cells were washed twice, incubated at 37°C for 3 min, fixed, immunostained with antibodies against Na/K-ATPase and imaged by confocal fluorescence microscopy. Scale bar, 10 µm. (**B**) Time-lapse fluorescence micrographs of wildtype (WT) or SMS-KO HeLa cells expressing GFP-tagged EqtSM or EqtSol and treated with 1500 U/ml SLO for the indicated time. Scale bar, 10 µm.

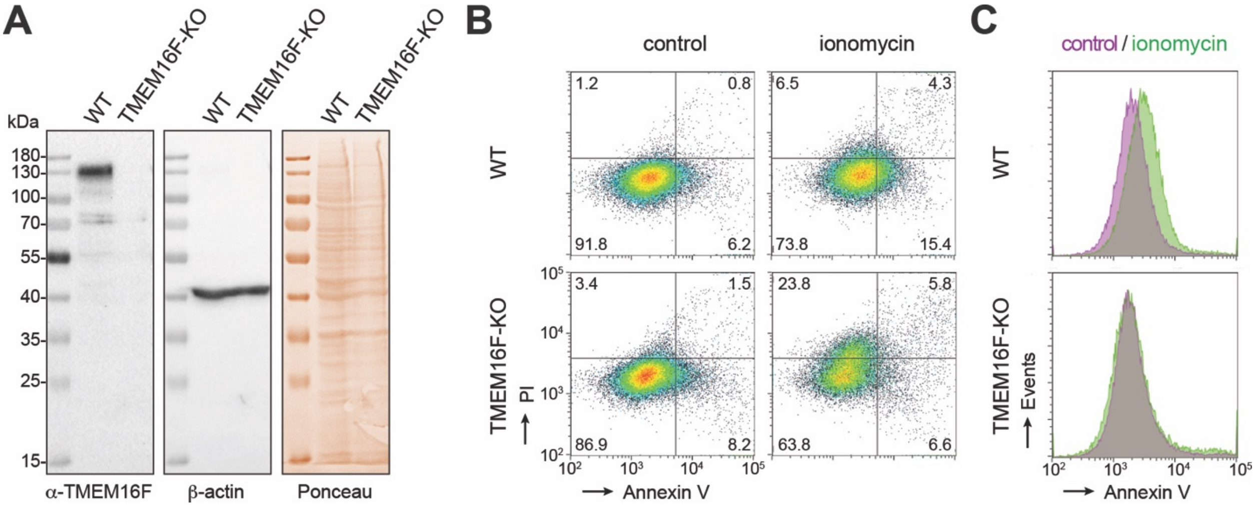


**Fig. S4. Characterization of TMEM16F-KO HeLa cells.**

(**A**) HeLa cells lacking TMEM16F (TMEM16F-KO) were created by CRISPR/Cas9. Loss of TMEM16F was confirmed by immunoblot analysis with antibodies against TMEM16F and β-actin. (**B**) Wildtype (WT) and TMEM16F-KO HeLa cells were incubated in the absence or presence of 15 μM ionomycin or 0.1% (v/v) DMSO for 10 min, stained with annexin V and propidium iodine, and then analyzed by flow cytometry. Representative dot plots are shown. (**C**) Histograms of annexin V staining of cells treated as in (B).

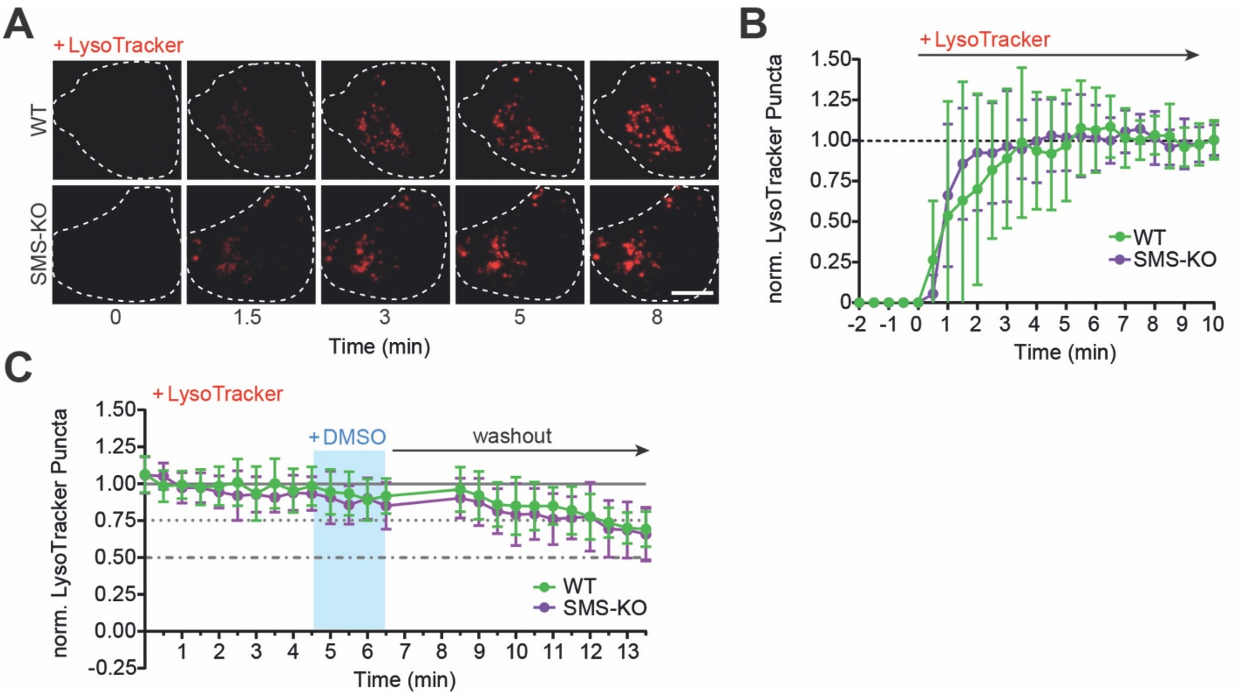


**Fig. S5. Wildtype and SMS-KO HeLa cells display similar LysoTracker labeling kinetics.**

(**A**) Time-lapse fluorescence micrographs of wildtype (WT) or SMS-KO HeLa cells incubated with 75 nM LysoTracker for the indicated time. Scale bar, 10 µm. (**B**) Time-course plotting LysoTracker-positive puncta in cells treated as in (B), normalized to the maximal number of puncta. Data are means ± SD from ≥ 14 cells per condition. (**C**) Time-course plotting LysoTracker-positive puncta in wildtype (WT) or SMS-KO HeLa cells during and after a 2 min-pulse of 0.06% (v/v) DMSO as vehicle control for GPN treatment. Scale bar, 10 µm.

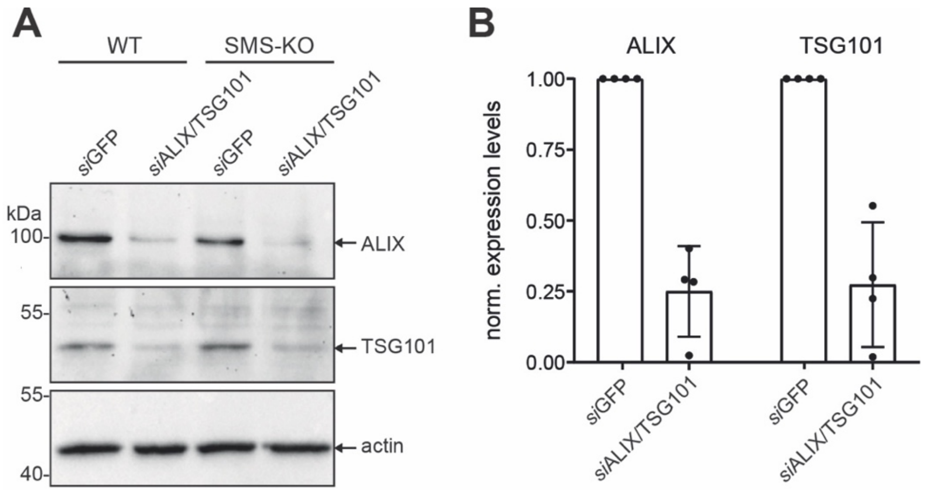


**Fig. S6. Validation of siRNA-mediated depletion of ALIX and TSG101.**

(**A**) Wildtype (WT) and SMS-KO HeLa cells were treated with siRNAs targeting GFP (siGFP) or ALIX and TSG101 (siALIX/TSG101) for 72 h, lysed, and then subjected to immunoblot analysis with antibodies against ALIX, TSG101 and β-actin. (**B**) Relative ALIX and TSG101 protein levels in cells treated as in (A) after normalization against actin levels. Data are means ± SD, $n = 4$.

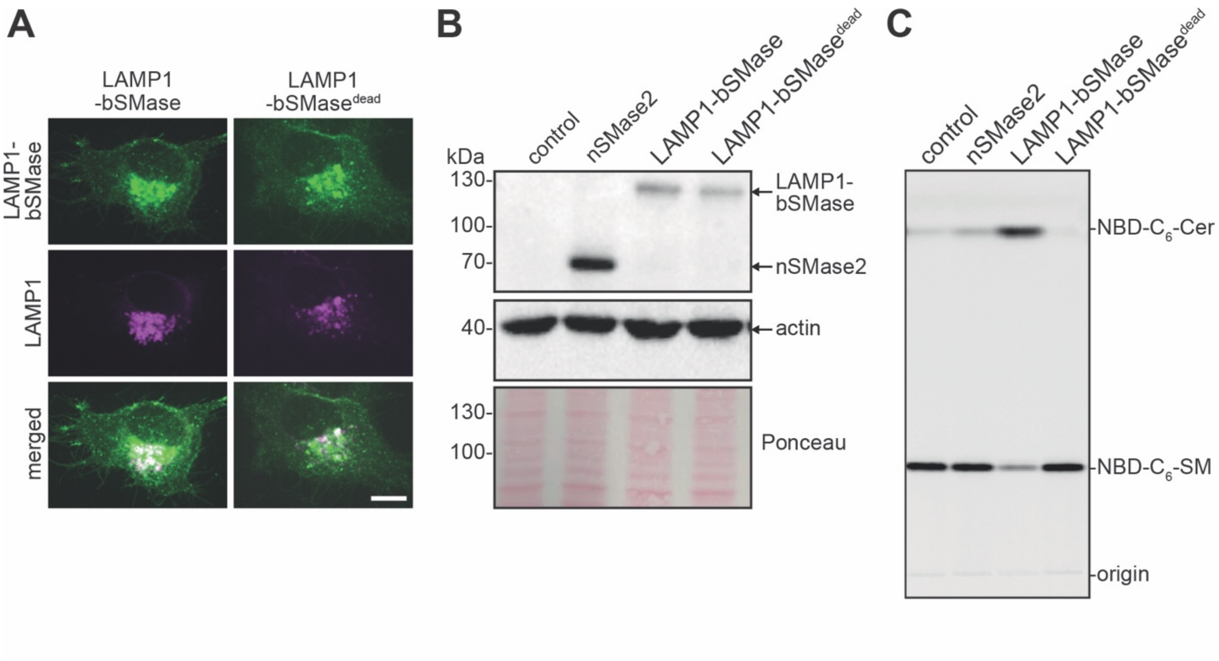


**Fig. S7. Characterization of LAMP1-bSMase fusion constructs.**

(**A**) HeLa cells co-transfected with mCherry-tagged LAMP1 (*magenta*) and GFP/V5-tagged LAMP1-bSMase or LAMP1-bSMase$^{dead}$ (*green*) were visualized by confocal fluorescence microscopy. Scale bar, 10 µm. (**B**) HeLa cells transfected with empty vector (control), V5-tagged nSMase2, GFP/V5-tagged LAMP1-bSMase or GFP/V5-tagged LAMP1-bSMase$^{dead}$ were lysed and subjected to immunoblot analysis using antibodies against V5 and β-actin. (**C**) TLC analysis of reaction products formed when lysates of cells treated as in (B) were incubated with 50 µM NBD-$C_6$-SM for 2 h at 37°C.

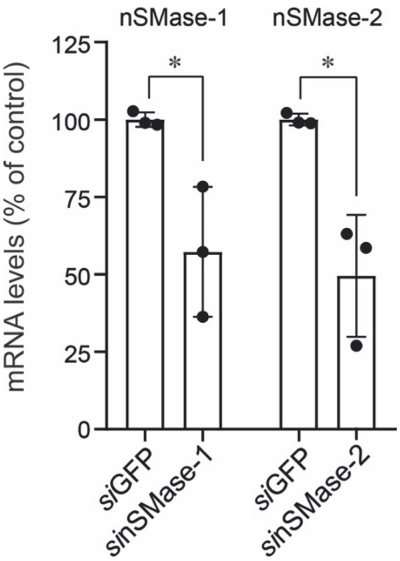


**Fig. S8. Validation of siRNA-mediated depletion of nSMase-1 and nSMase-2.**

HeLa cells were treated with siRNAs targeting GFP, nSMase-1 or nSMase-2 for 72 h. Total RNA was extracted, converted to cDNA and subjected to RT-qPCR using primers for actin, nSMase1 and nSMase2. Relative levels of nSMase1 and nSMase2 transcripts after normalization against actin are shown. Data are means ± SD, $n = 3$. Statistical significance was determined by unpaired two-tailed t-test.

**MOVIES**

**Movie S1.**

Time-lapse fluorescence images of HeLa cells co-expressing GFP-tagged EqtSM (left, *green*) and mCherry-tagged Gal3 (middle, *magenta*) treated with 1mM LLOMe for the indicated time. Images were captured every 30 s. Scale bar, 10 µm.

**Movie S2.**

Time-lapse fluorescence images of RAW264.7 cells expressing GFP-tagged EqtSM (*green*) infected with mCherry-expressing *M. marinum* (*magenta*). Images were captured every 60 s. Scale bar, 10 µm.

**Movie S3.**

Time-lapse fluorescence images of HeLa cells expressing GFP-tagged EqtSM locally wounded by a brief pulse from a 2-photon laser at high relative intensity (top wound, t = 0 s) and low relative intensity (bottom wound, t = 65 s). Images were captured every 5 s. Scale bar, 10 µm.

**Movie S4.**

Time-lapse fluorescence images of HeLa cells expressing GFP-tagged EqtSM (left, *green*) and labeled with LysoTracker (middle, *red*) during and after a 2 min-pulse of GPN (200 µM). Images were captured every 30 s. Scale bar, 10 µm.

**Movie S5.**

Time-lapse fluorescence images of HeLa cells co-expressing mKate-tagged EqtSM (left, *green*) and eGFP-tagged CHMP4B (middle, *magenta*) treated with 1 mM LLOMe for the indicated time. Images were captured every 10 s. Scale bar, 10 µm.

**Movie S6.**

Time-lapse fluorescence images of HeLa cells co-expressing mKate-tagged EqtSM (left, *green*) and GFP-tagged LAMP1-bSMase (middle, *magenta*) treated with 1 mM LLOMe for the indicated time. Images were captured every 30 s. Scale bar, 10 µm.

**Movie S7.**

Time-lapse images of HeLa cells co-expressing mKate-tagged EqtSM (left, *green*) and GFP-tagged LAMP1-bSMase$^{dead}$ (middle, *magenta*) treated with 1 mM LLOMe for the indicated time. Images were captured every 30 s. Scale bar, 10 μm.
